# Benchmark Averages Hide the Failures That Matter: Quantizing ESM-2 for Protein Variant-Effect Prediction

**DOI:** 10.64898/2026.08.10.744024

**Authors:** Qing Shao

## Abstract

We benchmark six numerical precision configurations for ESM-2 protein language models across throughput, memory footprint and predictive accuracy, on two workloads with sharply different characteristics: bulk embedding extraction and deep mutational scanning (DMS) variant-effect scoring. Accuracy is evaluated on the complete ProteinGym substitution benchmark — 201 assays, 2.41M variants — at **three model scales spanning 650M to 15B parameters**, with a paired bootstrap clustered on protein. Three findings follow, and each contradicts a common practice. First, **benchmark averages conceal the failure that decides deployability**: no configuration shifts mean correlation by more than 0.007 at any scale, yet INT8 dynamic quantization — indistinguishable from fp32 on that mean at 3B (*p* = 0.34) — takes a single assay from *ρ* = 0.591 to 0.223. Selection must be made on worst-case, not mean, behaviour. Second, **fidelity measured against fp32 bounds risk but cannot rank quality**: over 3015 assay/configuration pairs it predicts the *magnitude* of ground-truth change (*r* = 0.56–0.81) but not its *direction*, and the INT4 effect differs significantly between 650M and 3B (+0.0101, *p* = 0.0007) with no monotone trend to extrapolate. Third, **quantizing a large model is dominated by using a small one**: of eighteen scale/configuration combinations only three are Pareto-optimal over accuracy, memory and speed, and all three are 650M. The one catastrophic failure we observe is a defect of default *symmetric* activation scaling, not of W8A8 itself: asymmetric activation quantization, a one-line change needing no calibration, removes every damaged assay. We also give a label-free screen for at-risk targets, and report four measurement artifacts encountered during this study, three of which inverted the result they were meant to measure.

## 1 Introduction

Quantization is usually presented as a straightforward trade: reduce numerical precision, gain speed and memory, lose a little accuracy. This framing is inherited from large autoregressive language models, where it is broadly correct. We set out to test whether it transfers to ESM-2, a family of protein language models, and found that essentially every part of it requires qualification.

ESM-2 is a BERT-style bidirectional *encoder*. It performs a single forward pass with no KV cache and no autoregressive decode. This places it in a **compute-bound** regime rather than the memory-bandwidth-bound regime that governs LLM inference at batch size 1. The distinction is not academic: the dominant open-source quantization toolchains (GPTQ, AWQ, bitsandbytes NF4, GGUF) are all *weight-only*. They store INT4/INT8 weights and dequantize to bf16 immediately before an ordinary bf16 matrix multiply. When the bottleneck is memory bandwidth, this is a large win. When the bottleneck is arithmetic, as it is here, it adds work to an already-saturated kernel.

This paper addresses three questions:

1. What do the available quantization configurations actually cost and deliver on the three axes of speed, memory, and accuracy?
2. Does the answer depend on the workload? (It does, and the two workloads we study prefer opposite configurations.)
3. Do the fidelity metrics conventionally used to validate quantization predict real-world predictive accuracy? (Only partially, and in a way that matters.)

### Contributions

- An evaluation of six precision configurations for ESM-2 on the **complete ProteinGym substitution benchmark** — 201 assays, 2.41M variants, 40,775 masked positions — at **three model scales** (650M, 3B, 15B), rather than on drift from the full-precision model or on a handful of assays. 3618 assay/configuration runs in total.
- The finding that **benchmark-mean accuracy is nearly uninformative for deployment**. No configuration shifts mean correlation by more than 0.007 at any scale, while one that passes the mean test at *p* = 0.34 takes a single assay from *ρ* = 0.591 to 0.223. We argue that worst-case single-assay drift, not the mean, is the selection criterion, and give a label-free protocol for measuring it on a target (Algorithm 3).
- A quantitative characterisation of what drift-from-fp32 metrics buy. Over 3015 assay/configuration pairs they predict the magnitude of ground-truth change (*r* between 0.56 and 0.81) but not its direction. The direction does not merely vary — INT4 quantization is significantly harmful at 650M, significantly beneficial at 3B, and null at 15B, so there is no trend in scale to extrapolate either.
- The observation that **the configuration ranking is scale-dependent**. The two common ESM-2 workloads prefer opposite configurations at 650M and 3B, but FP8 dynamic quantization closes from 0.50× to 1.01× bf16 DMS throughput as scale grows, so the conflict dissolves by 15B. A recommendation derived at one scale does not transfer.
- Evidence that **scale does not buy variant-effect accuracy**: fp32 mean *ρ* falls monotonically 0.4459 → 0.4398 → 0.4323 across a 23-fold parameter increase, with no pairwise difference reaching significance.
- A documented account of **four measurement artifacts**, three of which inverted the quantity being measured while producing plausible, monotonic result tables — offered as reusable hazards rather than as local mistakes.
- The finding that the one catastrophic failure we observe is a **defect of default symmetric activation scaling, not of W8A8**: asymmetric activation quantization — one keyword, no calibration — removes all six damaged assays and improves worst-case drift from *ρ*_fp32_ = 0.393 to 0.965, and SmoothQuant does the same at higher cost. “Avoid INT8 dynamic quantization” would have been the wrong conclusion from the same benchmark.
- A **label-free screen** for which targets are at risk. The perturbation measured relative to the spread of the scores it perturbs separates damage cleanly across 3015 combinations: no combination above SNR = 8 loses more than 0.05 Spearman, and every collapse beyond 0.10 falls below 1.7. It needs one extra scoring pass and no experimental data.
- Evidence that the pipeline is **exactly reproducible**: 8 replicate runs on different GPUs give bitwise-identical scores over 19.3M comparisons, so the reported differences are properties of the configurations rather than of the runs.
- Open code, per-variant predictions for all 20 model/configuration combinations, and per-assay statistics (Section 9).

## 2 Background and Related Work

### Protein language models

ESM-2 [7] is a family of transformer encoders trained with masked language modelling on UniRef, following ESM-1b [12]. Architecturally these are BERT-style bidirectional encoders [2]: a single forward pass, no KV cache, no autoregressive decode. Meier et al. [9] established that such models predict variant effects zero-shot via *masked marginals* — masking a mutated position and comparing the log-probabilities of mutant and wild-type residues — which is the scoring protocol used throughout this work.

### Variant-effect benchmarks

ProteinGym [11] aggregates deep mutational scanning assays into a standard benchmark; the v1 substitution set used here contains 217 assays spanning 2.47M variants and five functional categories, including the mega-scale stability measurements of Tsuboyama et al. [15]. Reporting mean Spearman correlation across assays is the established summary statistic, and one of our contributions is to show what that statistic conceals.

### Post-training quantization

The dominant open-source toolchains are weight-only: GPTQ [6] and AWQ [8] store INT4 weights and dequantize before a bf16 matmul, while LLM.int8() [3] introduced mixed-precision decomposition to handle activation outliers. These were designed for autoregressive decoding, where batch-1 inference is memory-bandwidth-bound and dequantization is effectively free. Weight–activation (W8A8) schemes and the FP8 E4M3/E5M2 formats [10] instead reduce arithmetic cost, which is what a compute-bound encoder requires. We use torchao [14] for all configurations and PyTorch 2’s compiler stack [1], whose Inductor backend emits Triton [13] kernels; Section 4.1 shows that the compile mode, not the quantization scheme, determines whether W8A8 is viable at all. Quantization is also load-bearing for parameter-efficient fine-tuning [4], a setting we do not evaluate.

### Positioning

Quantization studies for protein language models are comparatively rare, and where accuracy is reported it is typically drift against the full-precision model on a small probe set. Our contribution is to replace that proxy with the full downstream benchmark at three model scales, and to show that the proxy is a valid risk bound but an invalid quality ranking.

## 3 Experimental Design

### 3.1 Configurations

Six configurations were evaluated, all applied via torchao to the encoder Linear layers only:

**Table 1:** Quantization configurations. Weight-only variants target memory; the dynamic (W8A8) variants also quantize activations and can therefore use low-precision tensor cores for the matrix multiply itself.

| Configuration | Class | Description |
| --- | --- | --- |
| fp32 | baseline | Reference for all fidelity measurements. |
| bf16 | baseline | The practical production default. |
| int8_wo | weight-only | INT8 weights, bf16 compute. Memory-oriented. |
| int4_wo | weight-only | INT4 weights, bf16 compute. Smallest footprint. |
| int8_dyn | W8A8 | INT8 weights *and* dynamically quantized activations; uses INT8 tensor cores. |
| fp8_dyn | W8A8 | FP8 (E4M3) weights and activations. Requires compute capability $\geq 8.9$; H200 only. |

### 3.2 Workloads

#### Bulk embedding extraction

Large length-bucketed batches under a token budget, measured in residues per second. This is the compute-bound regime.

#### DMS variant-effect scoring

The standard ESM masked-marginals protocol: for each mutated position, mask it and record log *p*(mut) − log *p*(wt) from the resulting distribution. Cost scales with the *number of mutated positions*, not with the number of variants, and the masked batch is narrow. This is the latency-bound regime.

Algorithm 1 states the scoring procedure as implemented. Two optimizations, both absent from typical reference implementations, are marked: only positions carrying a variant are masked (line 5), and the masked copies are batched rather than run one at a time (line 6). Together they are worth one to two orders of magnitude on a real assay — the median ProteinGym assay mutates 96 positions out of *L* = 222 — and they are applied identically to every configuration, so they change the absolute cost of the benchmark without affecting any comparison within it.

**Algorithm 1** Masked-marginals variant scoring, as benchmarked. Cost is 1|*P*| */B*l forward passes and is independent of |*V* |, so an assay with 16,000 variants over 900 positions costs the same as one with 900.

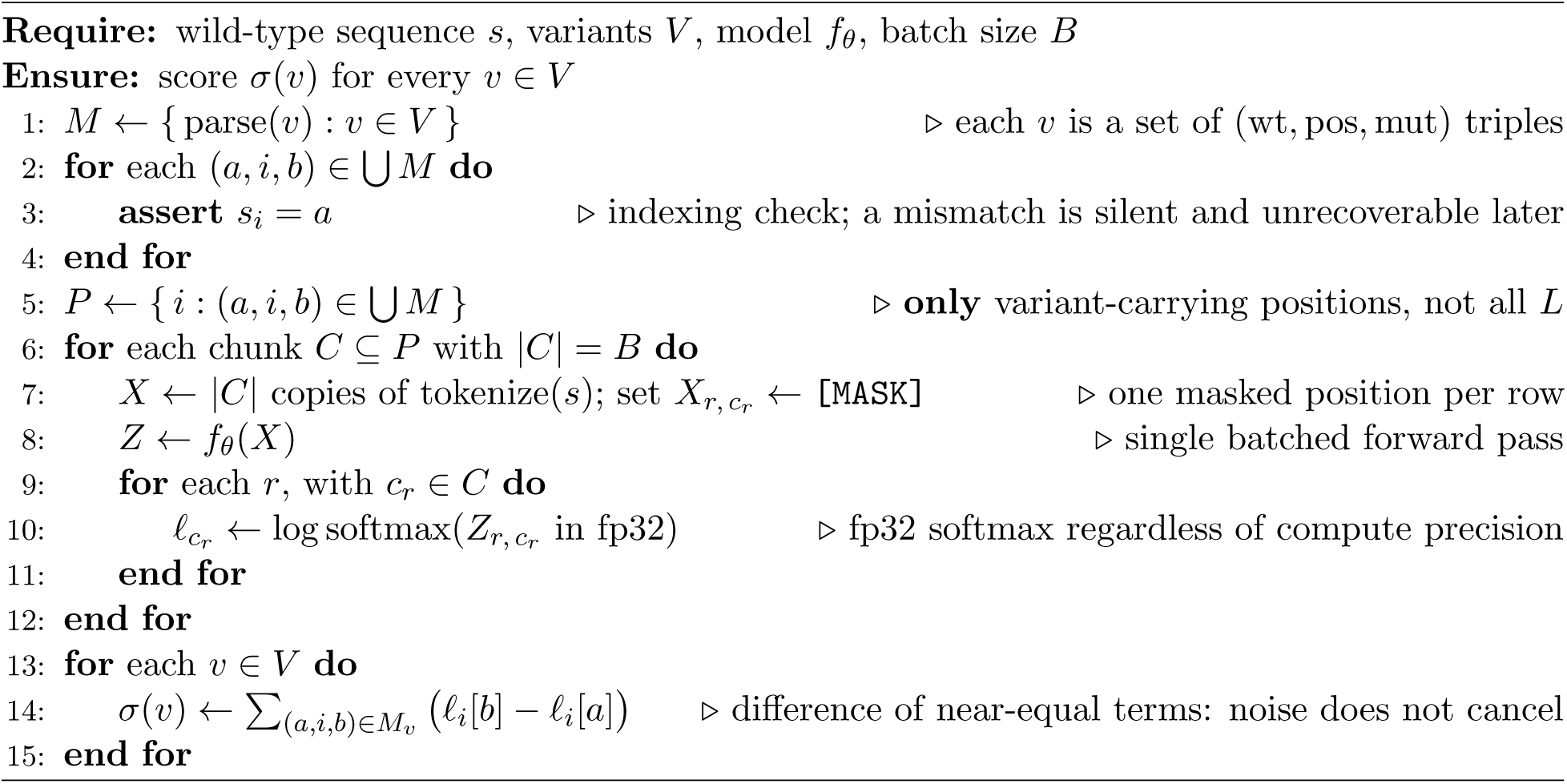

The final line is why DMS *ρ* is the most sensitive fidelity metric we record. The score is a difference of two nearly equal log-probabilities, so quantization error in *f_i_* does not cancel between the two terms; it is amplified relative to the signal.

### 3.3 Metrics

Three fidelity metrics were recorded, deliberately ordered by sensitivity:

1. **Embedding cosine similarity** (mean-pooled, excluding special tokens and padding). Forgiving: error averages out along the sequence.
2. **Logit KL divergence**, per residue. Moderately sensitive.
3. **DMS Spearman** *ρ* **against fp32.** Most sensitive, because the score is a *difference of two near-equal log-probabilities*, so quantization noise does not cancel and is amplified relative to signal.

All three are drift-from-fp32 measures. They quantify how far a configuration moves the answer, not whether it moves toward or away from the truth. To measure the latter, we added a fourth metric requiring real experimental data: **Spearman** *ρ* **against wet-lab DMS measurements**, from ProteinGym.

### 3.4 Real assay data

Three ProteinGym substitution assays were selected to span the sequence-length range, since peak memory during masked-marginals scoring grows with *L*: IF1 ECOLI Kelsic 2016 (*L* = 72), BLAT ECOLX Stiffler 2015 (*L* = 286) and HSP82 YEAST Flynn 2019 (*L* = 709), together 19,557 single-substitution variants over 1042 masked positions (Table 24). Every variant’s stated wild-type residue was verified against the reference target seq before use and all matched. This check is load-bearing — an off-by-one in position indexing does not raise an error, it silently scores the wrong positions and yields a plausible-looking correlation.

### 3.5 Full benchmark data

The three-assay analysis is superseded by a run over the complete ProteinGym substitution benchmark. All 217 assays were extracted from the ProteinGym v1 parquet release and every one passed wild-type validation across 2,465,767 variants. The 201 assays fitting ESM-2’s 1022-residue context were scored: 2,413,913 variants over 40,775 masked positions, spanning *L* = 37 to 934 and five function categories.

The masked-position count, not the variant count, sets the cost: masked-marginals scoring performs one forward pass per mutated position, so 2.41M variants are nearly free on top of 40,775 forwards.

An earlier three-assay path combined per-assay CSVs from the v0.1 release with wild-type sequences from the v1.0 reference file. The two agree on the 85 assays present in both and diverge elsewhere, including renames. That is acceptable for three hand-picked assays and unsafe at benchmark scale; the v1 parquet carries target seq inline, so scores and wild-type sequences come from a single release.

### 3.6 Comparability to published benchmark numbers

Two choices make our absolute correlations not directly comparable to a published ESM-2 ProteinGym figure, and we state both rather than leave a reader to discover them.

#### The aggregation convention matters more than any effect we measure

There is no single “the” ProteinGym average: an unweighted mean over assays, a mean of per-protein means, and a mean of per-function-category means are all defensible and all give different numbers. On our data they span 0.019 — roughly three times the largest quantization effect in the paper (Table 2).

**Table 2:** fp32 mean Spearman under four aggregation conventions. The flat mean we report is the most favourable of the four; a within-category mean is 0.019 lower at 3B. A reader comparing our numbers against a reference value must match the convention first. Crucially, the *differences between configurations* — everything this paper concludes from — are stable across all four (Table 3).

| Aggregation | ESM2-650M | ESM2-3B | ESM2-15B |
| --- | --- | --- | --- |
| Flat mean over 201 assays (this paper) | 0.4459 | 0.4398 | 0.4323 |
| Mean of per-protein means | 0.4403 | 0.4377 | 0.4321 |
| Mean of per-category means | 0.4297 | 0.4209 | 0.4152 |
| Post-stratified to the full 217 | 0.4406 | 0.4355 | 0.4290 |

**Table 3:** Change relative to fp32 at 3B under each convention. Signs are identical throughout and magnitudes agree to ∼0.001; the sole exception is int8 dyn under category weighting, which was not significant under any convention. The conclusions are therefore properties of the data rather than of the aggregation choice.

| Config | Flat | Per-protein | Per-category | Post-stratified |
| --- | --- | --- | --- | --- |
| `bf16` | +0.0002 | +0.0002 | +0.0002 | +0.0002 |
| `int8_wo` | +0.0001 | +0.0001 | +0.0004 | +0.0002 |
| `fp8_dyn` | +0.0005 | +0.0005 | +0.0012 | +0.0004 |
| `int8_dyn` | −0.0021 | −0.0027 | −0.0000 | −0.0022 |
| `int4_wo` | +0.0037 | +0.0032 | +0.0029 | +0.0034 |

**Table 4:**
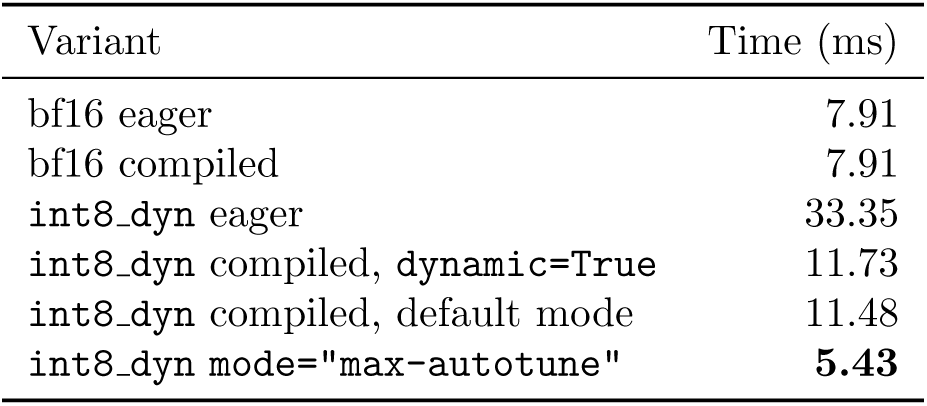
Only max-autotune allows Inductor to autotune the INT8 Triton matmuls, which is where the entire W8A8 benefit resides.

| Variant | Time (ms) |
| --- | --- |
| bf16 eager | 7.91 |
| bf16 compiled | 7.91 |
| int8_dyn eager | 33.35 |
| int8_dyn compiled, `dynamic=True` | 11.73 |
| int8_dyn compiled, default mode | 11.48 |
| int8_dyn mode="max-autotune" | **5.43** |

#### The 16 excluded assays are not a random sample

They exceed ESM-2’s context (*L* = 1154 to 3423) and account for 2.1% of benchmark variants, but their composition is skewed: **50% are low-MSA-depth against 13.9% of those retained**, and *none* are stability assays against 32.8% retained. Since low-MSA-depth assays score far worse (Table 15), excluding them inflates any unweighted mean. Post-stratifying the retained assays to the MSA-depth composition of all 217 gives 0.4406*/*0.4355*/*0.4290, i.e. the exclusion is worth roughly +0.004 at 3B. That is small against the reporting differences above but larger than several of the quantization effects, and it runs in the direction that flatters our absolute numbers.

The skew also has a second consequence worth flagging: because int4 wo costs most on low-MSA-depth assays at 650M (Table 15), excluding them *understates* its harm. Post-stratification moves that effect from −0.0064 to −0.0068.

### 3.7 Statistical treatment

Ground-truth correlations are compared using a **paired bootstrap**: *R* = 2000 resamples for a single assay, where the unit is the variant, and *R* = 20,000 for every benchmark-level result, where the unit is the assay and *R* = 2000 would floor the two-sided *p* at 0.001 — too coarse to separate the cases that matter in Section 4.7. Both the candidate configuration and the fp32 reference are scored on the *same* resample at each iteration, so shared noise cancels and the residual is attributable to the configuration. The pairing is not a refinement: for int4 wo the paired interval is 8.4× narrower than the unpaired one, because per-assay *ρ* spans 0.0–0.9 while the configuration effect is of order 0.006. Unpaired, every configuration reads as noise.

The *resampling unit* differs by scale of claim. For a single assay the unit is the variant, which establishes that a shift is not an artifact of the variant set. For the full benchmark the unit is the assay, since the claim becomes one about assays in general. Because the 201 assays cover only 173 distinct proteins, benchmark-level intervals use a **cluster bootstrap over proteins**: whole proteins are drawn and all their assays retained, so correlated replicates cannot count as independent evidence. Clustered and unclustered intervals agreed to the fourth decimal, which is reported as a measured result rather than assumed.

Intervals are percentile bootstrap [5]. They were checked against a normal approximation (d̅±1.96 SE) and agree to four decimals on every configuration, confirming the underlying distribution is near-symmetric and the percentile method is not distorting the interval. The procedure was validated on synthetic data with a known answer: a negligible perturbation was correctly classified as noise, a genuine improvement as real.

Algorithm 2 states the benchmark-level test. The pairing is line 5: one resample index is applied to the candidate and to the reference *together*. Without it the interval widens by 8.4× for int4 wo, because between-assay variance in *ρ* (spanning 0.0 to 0.9) is two orders of magnitude larger than the configuration effect (≈ 0.006), and every configuration is then reported as noise.

**Algorithm 2** Paired cluster bootstrap for Δ(mean *ρ*) against the fp32 reference. Setting G to singletons recovers the ordinary assay-level bootstrap; we report both.

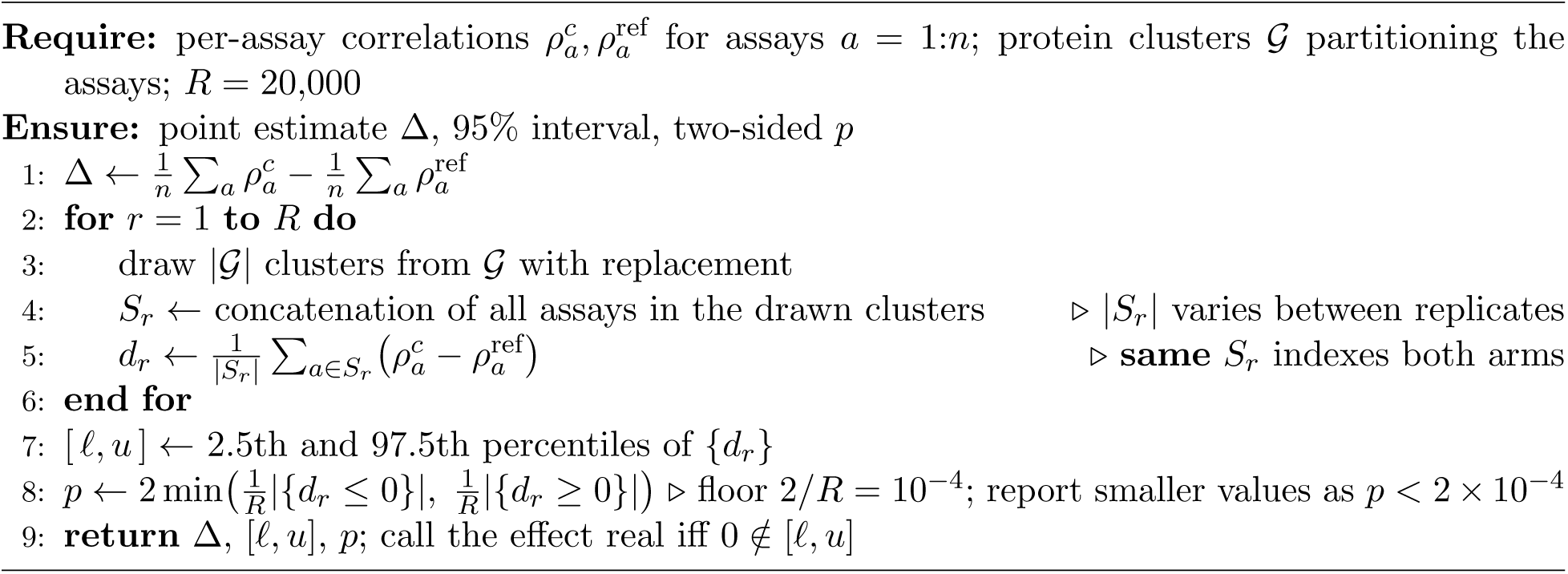

### 3.8 Measurement repeatability

Every difference we report is at 10^−4^ resolution or finer, and the tail statistic in Section 4.7.1 is a *maximum* over 201 assays — a quantity that any noise inflates, since the maximum of many noisy draws is large even when every true effect is zero. Neither number is interpretable without knowing what the pipeline does when nothing changes, so we measured it rather than assuming determinism.

The complete benchmark was re-run for all six configurations at 650M, and for bf16 and int8 dyn at 3B, as separate jobs on different physical GPUs. Across 8 model/configuration replicates and 19,311,304 per-variant score comparisons, **every score is identical** at the 10^−6^ precision to which scores are stored. Per-assay Spearman is unchanged to five decimals and benchmark means to six; the largest per-assay |Δ*ρ*|between two runs of the same configuration is 0.00000.

Three consequences follow.

- **The null for the tail statistic is exactly zero.** The maximum-over-201 objection does not apply here: there is no run-to-run variability for the maximum to select from, so a worst-assay |Δ*ρ*| of −0.369 is entirely attributable to the configuration.
- **Differences of** +0.0002 **are real differences.** They are properties of the configuration rather than of the run, which is what licenses reporting them at all.
- **The headline collapse reproduces exactly.** UBR5 HUMAN Tsuboyama 2023 1I2T gives *ρ* = 0.2226 under int8 dyn at 3B in both runs.

Three caveats. Determinism removes run-to-run variance and nothing else: the assay-sampling uncertainty that the bootstrap quantifies is untouched, and a difference being exactly reproducible does not make it large or important — a reproducible 0.0002 is still negligible. Both runs used H200 GPUs and one software stack, so determinism across GPU architectures or across torch/torchao versions is not established and should not be assumed. And scores are stored rounded to six decimal places, so “identical” means identical to 10^−6^, which is three orders of magnitude below the smallest effect discussed anywhere in this paper.

### 3.9 Hardware and software environment

Measurements were taken on an NVIDIA H200 (143 GB, sm 90, with FP8 tensor cores) under PyTorch 2.6.0+cu124 with SDPA attention, compiled with max-autotune-no-cudagraphs where compilation is used at all; an A100 (80 GB, sm 80, no FP8) was used where noted. Table 25 gives the full environment.

The cluster runs CentOS 7.6 with glibc 2.17 on every node, including the H200 nodes, which forced several choices. PyTorch moved to manylinux 2 28 at version 2.7, making **2.6.0 the newest installable release**. bitsandbytes GPU wheels require a newer glibc, so only the CPU-only 0.42.0 installs and its NF4 path is unavailable — 4-bit quantization therefore goes through torchao. The system GCC (4.8.5) cannot build Triton’s helper module, so torch.compile fails until CC/CXX are pointed at GCC 12.1.0. Finally, $HOME and /project are at quota, and the Triton, Inductor, HuggingFace, and pip caches all default there; all were relocated to /scratch. torchao nonetheless works because its INT8/FP8 paths dispatch to torch. int mm and torch. scaled mm, which live inside PyTorch and support sm 90.

## 4 Results

This section has four parts, and the first two disagree with each other. Sections 4.1–4.6 establish the speed, memory and fidelity behaviour of each configuration on the two workloads, taking ground truth from three hand-picked assays — enough to show that an effect exists on a particular assay, not enough to generalise from it. Section 4.7 repeats the accuracy measurement on the complete 201-assay benchmark at three model scales and overturns two conclusions the three-assay pilot supported; each retraction is stated where the original claim was made. The remainder diagnoses the one catastrophic failure the full benchmark exposes, repairs it, and derives a label-free screen that predicts which targets are at risk without experimental data. The last two subsections place all of it in a resource envelope and ask whether quantizing a large model is ever preferable to running a small one.

A reader interested only in the deployment conclusion can go to Table 8 (the per-assay tail, which is the selection criterion we argue for) and Table 19 (the cross-scale Pareto surface).

### 4.1 Compile mode determines whether W8A8 is viable at all

The most consequential configuration choice here is not the quantization scheme at all: it is the torch.compile mode. We measured a single ESM2-3B-shaped feed-forward block (2560 → 10240 → 2560) on an A100, at a batch of 32 × 512:

Measured in eager mode, W8A8 appears 4.2× *slower* than bf16 and would be discarded. Measured under max-autotune it is 1.46× faster. dynamic=True was avoided because it emits shape-generic kernels that surrender most of the gain; static shapes are kept manageable by padding to a fixed multiple of 128. torch. dynamo.explain confirms **zero graph breaks** for both bf16 and int8 dyn, establishing this as a kernel-selection effect rather than a Dynamo fallback.

### 4.2 Bulk embedding extraction

Three observations. First, the **fp32** → **bf16 step is worth** 8.4× — larger than everything quantization adds on top of it, and the single biggest lever available. Second, fp8 dyn dominates int8 dyn: it is faster (1.15× vs 1.05×) *and* dramatically more accurate (embedding cosine 0.999920 vs 0.989592; logit KL 4.6 × 10^−4^ vs 1.4 × 10^−2^), surrendering only 0.17 GB of peak memory. FP8’s E4M3 format has far greater dynamic range than INT8 at the same bit width, which matters because transformer activations contain outliers. Third, int4 wo is a trap for inference: 0.07× throughput and *worse* peak memory (7.69 GB) than fp8 dyn (7.24 GB) despite 40% smaller weights, because the dequantization path allocates transient buffers. Its value lies elsewhere, as a QLoRA base.

Separately, length-bucketed batching reduced padding waste from 25.0% (naive fixed-size batching) to **0.0%**. This is free and costs no accuracy — a larger effect than several of the quantization differences in the table.

### 4.3 DMS scoring: the opposite conclusion

bf16 is fastest at every assay size, and int8 dyn — the second-fastest configuration for bulk extraction — is up to 17× slower here. The masked batch (n positions × *L*) is too small to contain a matrix multiply large enough for low precision to pay for its quantize/dequantize overhead.

Notably, fp8 dyn closes on bf16 as the assay grows: 0.26×, 0.50×, 0.81× of bf16 throughput at *L* = 72, 286, 709 respectively. Its overhead is fixed per call while real work scales, so at *L* = 709 it costs 20% throughput for 43% less memory — a defensible trade that is indefensible at *L* = 72.

### 4.4 Memory during DMS scoring

Peak device memory during DMS scoring is the reverse of the bulk-extraction situation (Table 26). Masked-marginals scoring keeps the batch narrow, so the *O*(*L*^2^) attention term never dominates and weights remain roughly 90% of peak memory even at *L* = 709 (bf16: 5.29 GB of weights against 0.59 GB of activations). Quantization therefore addresses almost all of DMS peak memory, and int4 wo gives the smallest footprint at every length — 2.7× below bf16.

### 4.5 Ground-truth accuracy on ProteinGym

Spearman correlation against experimental measurement, with a 2000-sample paired bootstrap over variants, is reported for all three assays and six configurations at 3B in Table 27. Two findings emerge, and the second qualifies the first.

#### Quantization does not systematically degrade variant-effect accuracy

Of the 15 as-say/configuration pairs, 11 shifts are upward. Most are statistically indistinguishable from zero. Strikingly, int4 wo — the worst configuration on every fidelity metric — produces the *best* agreement with experiment on BLAT (+0.0495, CI [+0.0433, +0.0561]). This survives the bootstrap and is not a resampling artifact.

#### Fidelity-vs-fp32 predicts the magnitude of the shift, not its sign

Across all 15 pairs, the correlation between fidelity loss (1 − *ρ*_fp32_) and the absolute ground-truth shift is *r* = 0.87 (Pearson), *ρ* = 0.76 (Spearman). But int4 wo moves +0.0495 on BLAT and −0.0167 on IF1. The direction is a property of the assay, not of the configuration, and is not predictable from any drift-from-fp32 measurement.

The practical consequence is that bf16 and int8 wo shift ground-truth agreement by at most 0.002 on any assay tested — below anything an experiment could resolve — and are therefore safe. int4 wo is a ±0.05 gamble on an unknown sign.

This supersedes an earlier conclusion drawn from a synthetic 40-residue mutational scan, which ranked int8 dyn (*ρ*_fp32_ = 0.9928) as risky and fp8 dyn (0.9988) as clearly safer. Both numbers were correct; neither predicted experimental accuracy. On BLAT the “risky” configuration gained +0.027.

We note that the assay itself matters far more than any quantization decision: *ρ*_expt_ ranges from 0.29 (HSP82) to 0.59 (BLAT) at 3B. Choosing which assay to trust is a 2× effect; choosing a quantization configuration is a 1% effect.

### 4.6 Model-scale consistency

The same three assays were run on ESM2-650M. The qualitative picture holds: bf16 fastest, int8 dyn slowest, ground-truth shifts small — the largest 650M shift was −0.0063 (int8 dyn on IF1) against *ρ*_fp32_ of 0.9954. The larger and more clearly significant shifts at 3B suggested that sensitivity to quantization grows with model size.

The pilot appeared to say something stronger as well. The fp32 650M model correlated with experiment *better* than fp32 3B on two of the three assays, and on BLAT by 0.1422 (0.7315 against 0.5893) — roughly *three times* the largest effect any quantization configuration produced in this study. The inference drawn was that model selection dominates precision selection by a wide margin.

*That inference does not survive Section 4.7*. Over 201 assays the two models are statistically indistinguishable (*p* = 0.35), and adding a third scale makes the picture worse for the larger checkpoint rather than better: fp32 mean *ρ* falls monotonically 0.4459 → 0.4398 → 0.4323 from 650M to 3B to 15B, with no pairwise difference significant (15B − 650M = −0.0137, *p* = 0.14). The BLAT gap is real for BLAT and does not generalise — the same error, made the same way, as the int4 wo result three subsections above. What survives is only the negative claim: there is no evidence that scale buys variant-effect accuracy, and 15B costs 5.4× the compute of 3B for the lowest mean of the three.

### 4.7 The full benchmark at three model scales

Every accuracy conclusion above rests on three assays and one model. Three assays are enough to establish that an effect exists on a particular assay and not enough to establish anything about assays in general, because the unit that varies — the assay — is held fixed. One model scale is enough to establish that an effect exists for that checkpoint and not enough to know whether it transfers. The whole ProteinGym substitution benchmark was therefore run at three scales: all 217 assays validated, the 201 within ESM-2’s 1022-residue context scored, 2,413,913 variants over 40,775 masked positions, six configurations, at 650M, 3B and 15B parameters. 3618 assay/configuration runs, 16.2 GPU-hours.

Two changes to the statistical treatment follow from the larger sample. The resampling unit becomes the **assay** rather than the variant, since the question is now whether a configuration degrades prediction in general. And because the 201 assays cover only 173 distinct proteins — BLAT ECOLX appears four times — the reported interval is a **cluster bootstrap over proteins** (Algorithm 2). The two intervals agree to the fourth decimal, so the clustering was immaterial here; it is reported because that had to be measured rather than assumed.

#### 4.7.1 The benchmark mean is the wrong safety criterion

Every Δ in Table 7 is below 0.007. The per-assay tail is not (Figure 1, Table 8). int8 dyn at 3B is indistinguishable from fp32 on the benchmark mean (Δ = −0.0021, *p* = 0.34) while destroying UBR5 HUMAN Tsuboyama 2023 1I2T, where correlation with experiment falls from 0.591 to 0.223 at *ρ*_fp32_ = 0.393. A configuration can pass the benchmark average and be unusable on the one protein a given project cares about. This is the practical result of the whole study: benchmark-mean significance testing answers a question about populations, and deployment is a question about a specific instance.

**Figure 1:**
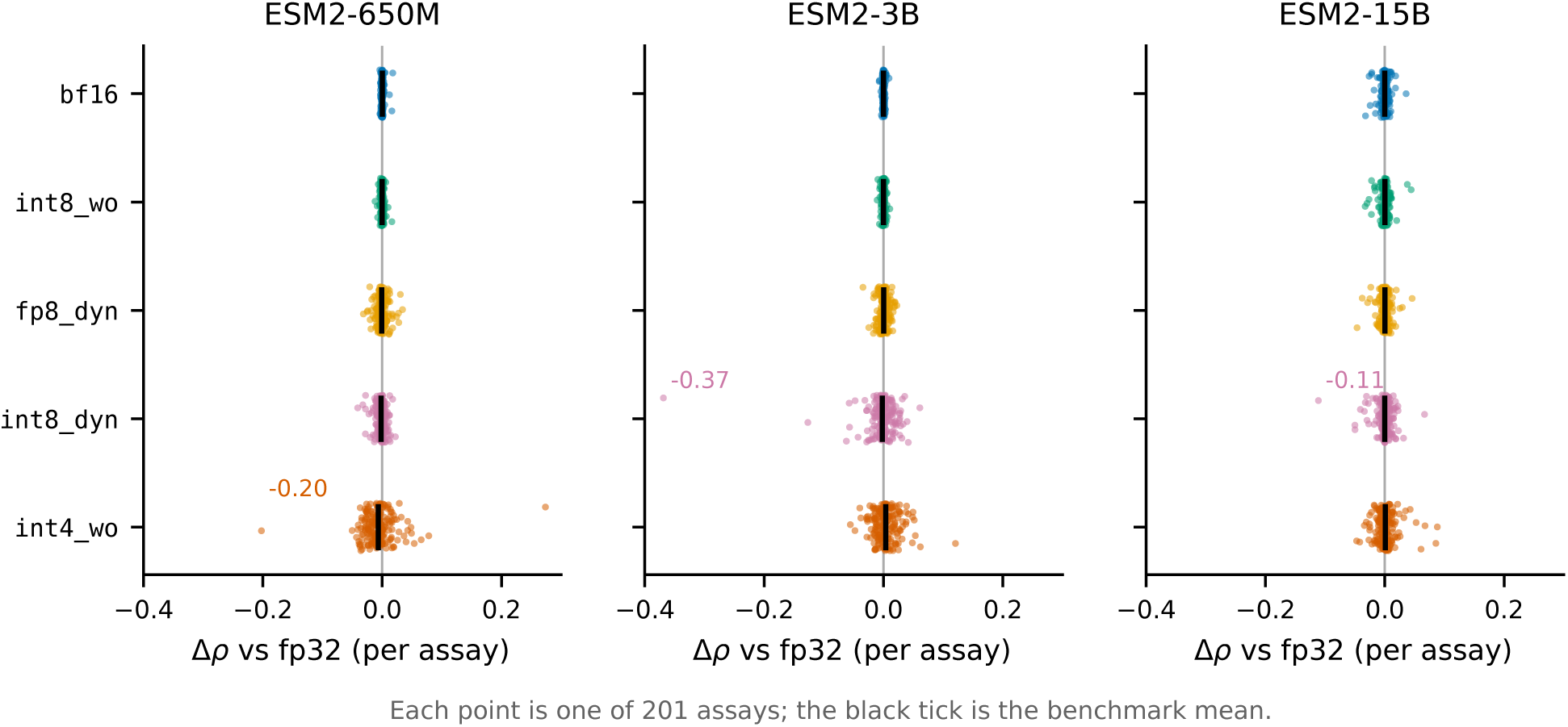
Per-assay change in ground-truth correlation for each configuration, across all 201 ProteinGym assays and all three model scales. Black ticks mark the benchmark mean, which is within 0.007 of zero in every panel. bf16 and int8 wo are tight at every assay and every scale; int8 dyn and int4 wo have means that are equally unremarkable and tails that reach −0.37 and −0.20. Selection on the mean cannot distinguish these cases; selection on the tail separates them immediately.

**Table 5:** ESM2-3B, H200, compiled with max-autotune-no-cudagraphs, SDPA attention, 64 sequences of length 512–1024 under a 16,384-token budget. Reproduced across five independent jobs: bf16 mean 56,789 residues/s, range 55,941–57,422 (±1.3%); fp32 mean 6,752, range 6,687–6,814. The int8 dyn asym row comes from the fifth of those jobs, run at identical settings; its bf16 (57,377 residues/s) and int8 dyn (60,023) fall inside the ranges above, so the rows are comparable.

| Config | Weights (GB) | Peak (GB) | Residues/s | vs bf16 | Emb. cos | Logit KL | DMS $\rho_{fp32}$ |
| --- | --- | --- | --- | --- | --- | --- | --- |
| fp32 (ref) | 10.72 | 25.37 | 6,750 | $0.12\times$ | — | — | — |
| bf16 | 5.29 | 12.63 | 57,057 | $1.00\times$ | 0.999949 | $1.17e-04$ | 0.9998 |
| int8_wo | 2.66 | 8.48 | 54,816 | $0.96\times$ | 0.999959 | $1.05e-04$ | 0.9998 |
| int8_dyn | 2.65 | **7.07** | 59,996 | $1.05\times$ | 0.989592 | $1.40e-02$ | 0.9928 |
| int8_dyn_asym | 2.65 | 7.26 | 51,383 | $0.90\times$ | 0.999469 | $1.41e-03$ | 0.9972 |
| fp8_dyn | 2.65 | 7.24 | **65,376** | **$1.15\times$** | 0.999920 | $4.60e-04$ | 0.9988 |
| int4_wo | **1.61** | 7.69 | 3,813 | $0.07\times$ | 0.998904 | $1.89e-03$ | 0.9975 |

**Table 6:** DMS scoring throughput, ESM2-3B, uncompiled. bf16 wins at every problem size; quantization is pure overhead in this regime.

| Config | Variants/s |  |  |
| --- | --- | --- | --- |
| | IF1 ($L=72$) | BLAT ($L=286$) | HSP82 ($L=709$) |
| `bf16` | **6,979** | **3,573** | **1,357** |
| `int8_wo` | 4,830 | 2,511 | 1,066 |
| `fp8_dyn` | 1,813 | 1,796 | 1,103 |
| `fp32` | 1,460 | 447 | 187 |
| `int8_dyn` | 410 | 382 | 299 |
| `int4_wo` | 1,045 | 282 | 112 |

**Table 7:** Change in mean Spearman against experiment relative to fp32, over 201 assays, at each model scale. Bold marks intervals excluding zero under the protein-clustered paired bootstrap at *R* = 20,000 (Algorithm 2). int4 wo is significant in *opposite directions* at 650M and 3B and null at 15B; the effect does not merely vary in size, it has no consistent sign and no monotone trend in scale. fp32 mean *ρ* is 0.4459, 0.4398 and 0.4323 respectively.

| Config | ESM2-650M |  | ESM2-3B |  | ESM2-15B |  |
| --- | --- | --- | --- | --- | --- | --- |
| | $\Delta$ | 95% CI | $\Delta$ | 95% CI | $\Delta$ | 95% CI |
| `bf16` | +0.0000 | $[-0.0003, +0.0004]$ | +0.0002 | $[-0.0001, +0.0004]$ | +0.0000 | $[-0.0009, +0.0011]$ |
| `int8_wo` | -0.0002 | $[-0.0006, +0.0002]$ | +0.0001 | $[-0.0002, +0.0005]$ | +0.0002 | $[-0.0009, +0.0013]$ |
| `fp8_dyn` | -0.0007 | $[-0.0020, +0.0005]$ | +0.0005 | $[-0.0007, +0.0018]$ | +0.0002 | $[-0.0010, +0.0015]$ |
| `int8_dyn` | **-0.0020** | $[-0.0031, -0.0009]$ | -0.0021 | $[-0.0072, +0.0017]$ | -0.0000 | $[-0.0022, +0.0020]$ |
| `int4_wo` | **-0.0064** | $[-0.0107, -0.0017]$ | **+0.0037** | $[+0.0008, +0.0069]$ | +0.0009 | $[-0.0014, +0.0034]$ |

**Table 8:** Worst single-assay change in ground-truth correlation, with the count of assays moving by more than 0.05 in parentheses, out of 201. The catastrophic tail does not sit with a fixed configuration: it is int4 wo at 650M and int8 dyn at 3B.

| Config | ESM2-650M | ESM2-3B | ESM2-15B |
| --- | --- | --- | --- |
| `bf16` | −0.0040 (0) | −0.0072 (0) | −0.0323 (0) |
| `int8_wo` | −0.0120 (0) | −0.0106 (0) | −0.0329 (0) |
| `fp8_dyn` | −0.0316 (0) | −0.0344 (0) | −0.0465 (0) |
| `int8_dyn` | −0.0411 (0) | **−0.3688** (6) | −0.1112 (3) |
| `int4_wo` | **−0.2024** (6) | −0.0556 (5) | −0.0472 (5) |

Table 8 adds a second warning. The catastrophic tail is not a fixed property of a configuration either — it belongs to int4 wo at 650M and to int8 dyn at 3B. Knowing which configuration was dangerous at one scale does not tell you which will be dangerous at another.

##### Conditioning the tail on the reference having signal

A large drop in rank correlation on an assay where fp32 itself scores near zero is rank instability, not damage: the configuration was useless on that target either way, and counting such cases inflates the tail. Restricting to the 156–157 assays (of 201) where fp32 reaches *ρ >* 0.3 separates the two (Table 9).

**Table 9:** Worst single-assay Δ*ρ* unconditioned, and restricted to assays where the fp32 reference itself achieves *ρ >* 0.3. The 3B int8 dyn collapse is unchanged by conditioning — fp32 reaches 0.591 on that assay — while the 650M int4 wo and 15B int8 dyn figures shrink substantially, because they occur on assays where fp32 reaches only 0.270 and 0.290. Counts in parentheses are assays with |Δ*ρ*| *>* 0.05 among those retained.

| Config | worst $\Delta\rho$, all 201 | | | worst $\Delta\rho$, $\rho_{\text{fp32}} > 0.3$ | | |
| --- | --- | --- | --- | --- | --- | --- |
|  | 650M | 3B | 15B | 650M | 3B | 15B |
| `bf16` | −0.004 | −0.007 | −0.032 | −0.004 (0) | −0.007 (0) | −0.022 (0) |
| `int8_wo` | −0.012 | −0.011 | −0.033 | −0.012 (0) | −0.009 (0) | −0.018 (0) |
| `fp8_dyn` | −0.032 | −0.034 | −0.046 | −0.025 (0) | −0.034 (0) | −0.025 (0) |
| `int8_dyn` | −0.041 | **−0.369** | −0.111 | −0.032 (0) | **−0.369** (2) | −0.030 (0) |
| `int4_wo` | −0.202 | −0.056 | −0.047 | −0.050 (4) | −0.056 (5) | −0.047 (2) |

The threshold is a judgement call, and we report its sensitivity rather than let one cut carry the conclusion. The 3B int8 dyn collapse is −0.369 at every cut from 0.0 to 0.4; the 650M int4 wo figure is −0.202 up to a cut of 0.2 and −0.050 beyond it. Our central example is therefore robust and the −0.20 figure is not, which is why the recommendation table now cites −0.06 rather than −0.20 as the drift a practitioner should plan for.

Conditioning also sharpens the practical statement. On assays where the model is usable at all, int8 dyn at 3B damages 2 of 157 and int4 wo damages 4–5 of 157 by more than 0.05: roughly a 1–3% chance that a given target is affected. That is the number to act on, and it is not visible in a mean.

#### 4.7.2 At 15B the safe configurations degrade and the aggressive one improves

Scale does not move all configurations in the same direction, and the crossover is visible only with a third point (Figure 2c, Table 10). bf16 and int8 wo do not drift measurably from fp32 on *any* of the 402 assay/model pairs at 650M and 3B, and then drift on 13 and 15 assays respectively at 15B, with worst-case rank correlation falling to 0.857 and 0.855. int4 wo moves the other way: 182 assays below *ρ*_fp32_ = 0.99 at 650M, 154 at 3B, 82 at 15B.

**Figure 2:**
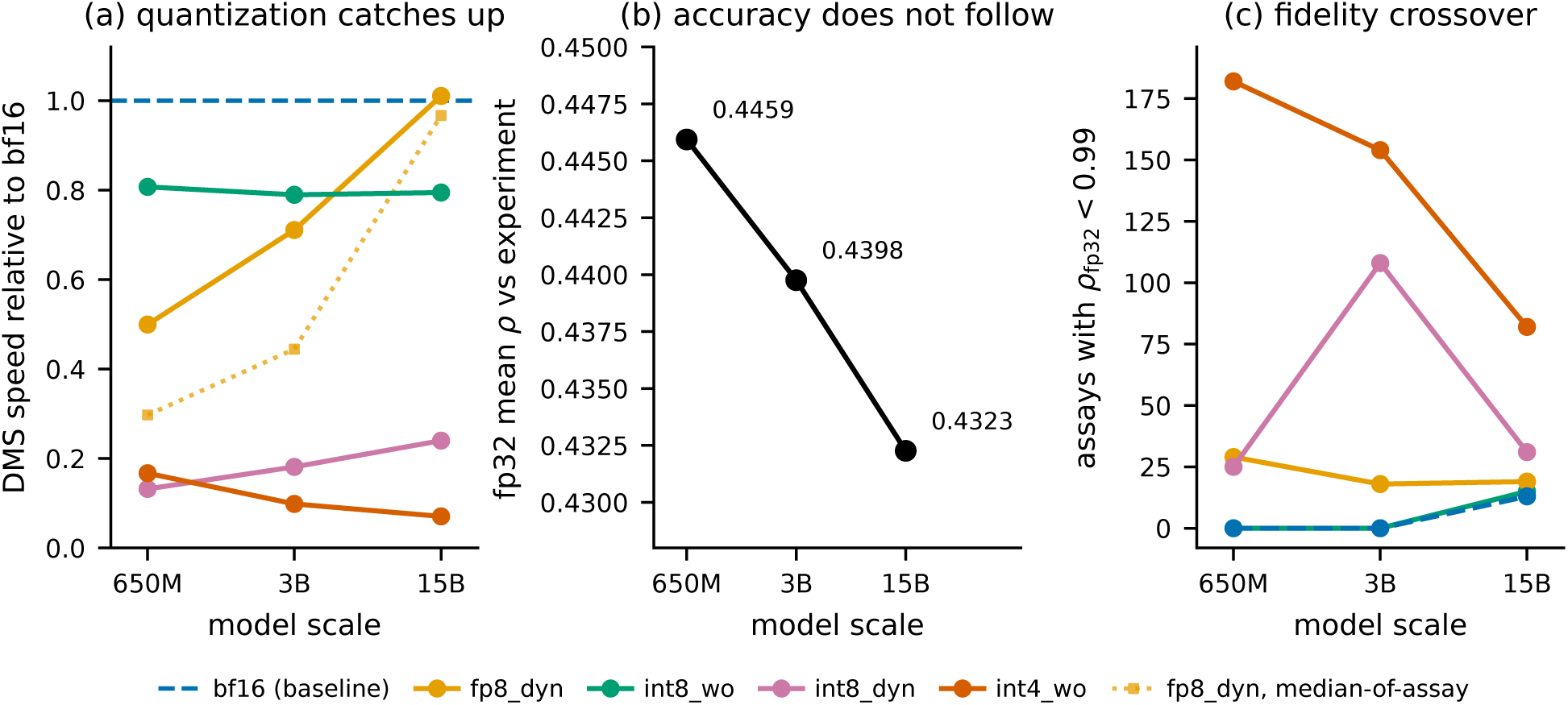
Three effects visible only across scale. (a) DMS speed relative to bf16, as aggregate throughput: fp8 dyn converges on bf16 while the weight-only and INT4 paths do not, so “quantization is pure overhead for DMS” is scale-bounded. The dotted line repeats fp8 dyn under the median-of-assay convention — the only configuration for which the choice is visible, because its overhead is fixed per call and therefore worst on the short assays a median weights most heavily. The convergence holds either way. (b) fp32 mean Spearman against experiment falls monotonically across a 23-fold parameter increase; no pairwise difference is significant, but there is no evidence that scale buys variant-effect accuracy, and 15B costs 5.4× the compute of 3B for the lowest of the three. (c) Number of assays where a configuration drifts from fp32 (*ρ*_fp32_ *<* 0.99): bf16 and int8 wo acquire a tail only at 15B, while int4 wo sheds one. The safe and the aggressive configurations move in opposite directions.

**Table 10:** Fidelity against fp32 by scale: number of assays with *ρ*_fp32_ *<* 0.99 (of 201), and the worst single-assay value. The two columns move in opposite directions with scale — the configurations that are safe at small scale are the ones that acquire a tail at 15B, while the most aggressive one loses its tail. Medians stay above 0.99 everywhere, so none of this is visible in a summary statistic.

| Config | assays with $\rho_{\text{fp32}} < 0.99$ | | | worst $\rho_{\text{fp32}}$ | | |
| --- | --- | --- | --- | --- | --- | --- |
|  | 650M | 3B | 15B | 650M | 3B | 15B |
| `bf16` | 0 | 0 | 13 | 0.9974 | 0.9905 | 0.8569 |
| `int8_wo` | 0 | 0 | 15 | 0.9963 | 0.9968 | 0.8548 |
| `fp8_dyn` | 29 | 18 | 19 | 0.9556 | 0.9770 | 0.8389 |
| `int8_dyn` | 25 | 108 | 31 | 0.8894 | 0.3930 | 0.7645 |
| `int4_wo` | 182 | 154 | 82 | 0.6707 | 0.8791 | 0.8505 |

**Table 11:**
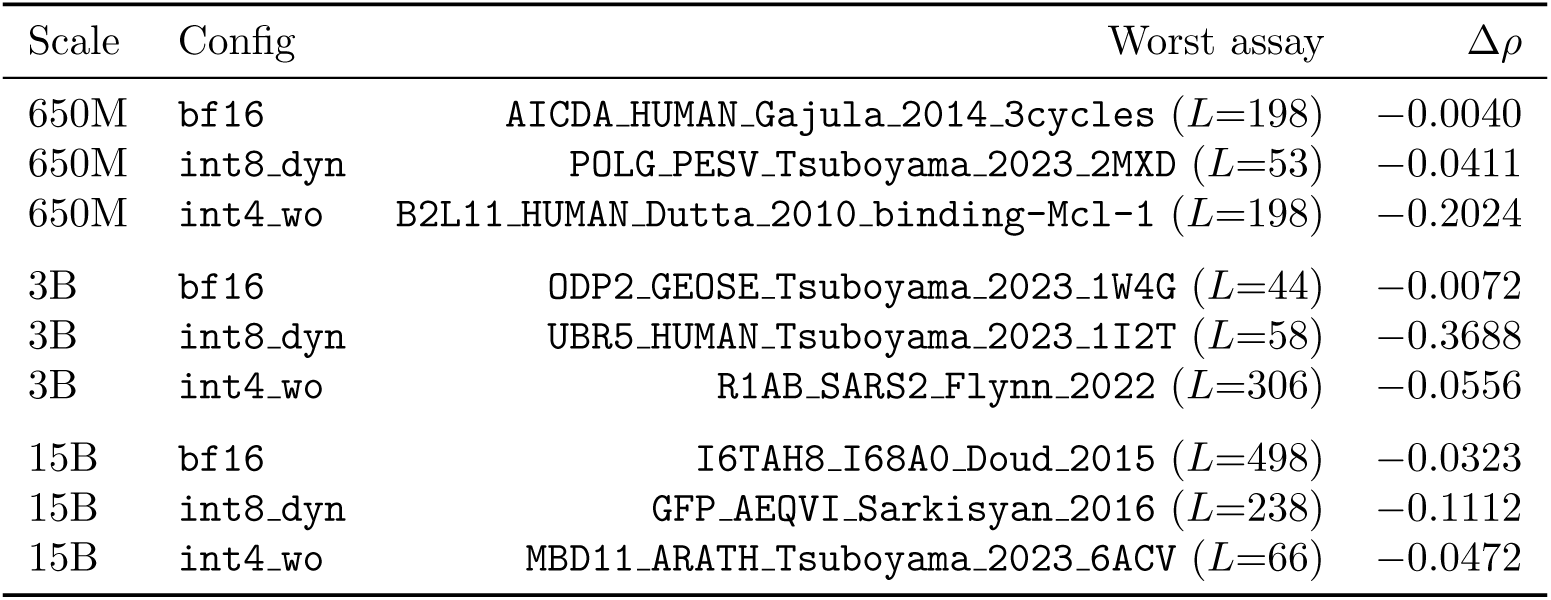
The single worst-affected assay, by configuration and scale. No assay appears twice. Identifying the vulnerable target at one scale does not identify it at another, which is why the screen in Algorithm 3 has to be run against the model actually being deployed.

A natural reading is that a larger model carries more redundancy, so removing weight precision costs proportionally less, while its activation dynamic range grows until bf16’s exponent becomes the binding constraint. We do not have the evidence to assert that mechanism: it would need a per-layer error analysis and an fp16-versus-bf16 comparison, neither of which we ran. What the data supports is narrower and still consequential — **the identity of the risky configuration is not stable across scale**, so a configuration cleared at one scale has not thereby been cleared at another.

The 15B bf16 degradation is not a long-sequence artifact. Spearman correlation between assay length and *ρ*_fp32_ is −0.14, and the five worst assays span *L* = 222 to 861.

#### 4.7.3 Resource envelope at 15B

Peak memory at 15B is almost entirely weights, and the practical consequence is a hardware boundary rather than a preference: fp32 needs 57 GB of weights before activations, so the reference configuration alone does not fit an 80 GB A100, and bf16 at 28.6 GB leaves little room beside it. Every quantized configuration returns the model to a single accelerator with margin, which at this scale is what quantization is for — the accuracy question of Table 7 only arises once the model runs at all (Table 12).

**Table 12:** ESM2-15B footprint. Peak memory during masked-marginals scoring is almost entirely weights: the spread across 201 assays spanning *L* = 37 to 934 is under 3 GB for every configuration. Quantization therefore addresses essentially all of the DMS memory cost at this scale, and fp32 alone rules out an 80 GB A100.

| Config | Weights (GB) | Peak memory over 201 assays (GB) |  |  |
| --- | --- | --- | --- | --- |
|  |  | min | median | max |
| fp32 | 56.36 | 56.51 | 57.09 | 59.30 |
| bf16 | 28.18 | 28.27 | 28.57 | 29.69 |
| int8_wo | 14.14 | 14.39 | 14.53 | 15.64 |
| fp8_dyn | 14.12 | 14.22 | 14.57 | 15.91 |
| int8_dyn | 14.12 | 14.51 | 14.90 | 16.06 |
| int4_wo | 7.53 | 7.62 | 7.91 | 9.03 |

#### 4.7.4 The configuration ranking is scale-dependent

Section 6.1 reports that the two workloads prefer opposite configurations, on the evidence that fp8 dyn wins bulk extraction and loses badly at DMS scoring. That holds at 650M and 3B and stops holding at 15B.

At 15B, fp8 dyn scores the benchmark in 0.381 GPU-hours against bf16’s 0.385, with peak memory 14.57 against 28.57 GB and no measurable accuracy cost. It is simply the better choice, which it is not at either smaller scale. The workload conflict is therefore a small-model phenomenon, and a recommendation derived at 650M or 3B does not transfer upward (Figure 2).

##### Which speed convention, and why it matters here

Section 3.6 argues that an aggregation convention must be stated because the choice can exceed the effect being reported. That applies to speed at least as sharply as to accuracy. Averaging the 201 per-assay rates answers “how much slower on a typical assay”; dividing total positions by total seconds answers “how much longer does the job take”. For four of the five configurations the two agree to within 0.10, and for fp8 dyn they differ by up to 1.7× — 0.50 against 0.30 at 650M. The gap is not noise but the mechanism restated: fp8 dyn’s per-assay ratio at 3B rises from 0.27 on the shortest quartile of assays to 0.77 on the longest, so an assay-weighted summary is dominated by the short assays where its fixed cost is worst, and a position-weighted one by the long assays where it is amortised.

We report throughput as the primary figure, because it is what a user experiences and because it is the convention already implied by the wall-clock column of Table 20. The qualitative conclusion — that the penalty shrinks monotonically with scale and vanishes at 15B — holds under both. A third convention, the unweighted *mean* of per-assay rates, gives 0.31, 0.38 and 0.56; we do not adopt it, but note that it is the one convention under which “the conflict dissolves at 15B” would be an overstatement, and that we would not have known which we had used had we not computed all three.

#### 4.7.5 What the larger sample changed

##### The three-assay verdict was backwards, and the correction does not resolve into a trend

Section 4.5 concluded that quantization does not systematically degrade variant-effect accuracy, on the evidence that 11 of 15 shifts were upward and that int4 wo gained +0.0495 on BLAT. At benchmark scale int4 wo moves −0.0064 at 650M, +0.0037 at 3B and +0.0009 at 15B. Stating this as “significant in opposite directions” would rest on two separate tests against fp32 whose verdicts are then compared by eye, and those tests are only nominally significant: Table 7 reports 15 comparisons (five configurations at three scales), against which a Bonferroni threshold is 0.0033, and the two int4 wo results give *p* = 0.0102 and *p* = 0.0104. Neither survives.

The claim we actually want is that the effect *differs* between scales, and that is one paired test on the same 201 assays rather than two tests and a comparison. It is also the stronger test (Table 14):

**Table 13:** DMS scoring speed relative to bf16 over the full 201-assay benchmark, under both aggregation conventions. *Throughput* is total masked positions over total seconds, and is the figure consistent with the wall-clock hours of Table 20; *median* is the median of the 201 per-assay rates, which weights a 40-position assay as heavily as a 900-position one. bf16 runs at 174.2, 93.7 and 29.4 positions/s by throughput and 347.7, 200.6 and 56.7 by median. Weight-only int8 wo holds a constant ratio under either convention, as expected for a scheme adding fixed dequantization work per weight. fp8 dyn closes on bf16 as scale grows under both, reaching parity at 15B: its overhead is per-call while the useful work grows with hidden size, so by 15B the GEMMs are large enough to be compute-bound even in the narrow-batch DMS regime. That same per-call overhead is why the two conventions separate for fp8 dyn and for no other configuration — a fixed cost per forward pass is proportionally worst on short assays, which the median weights most heavily. The paragraph below the table sets out the choice.

| Config | ESM2-650M |  | ESM2-3B |  | ESM2-15B |  |
| --- | --- | --- | --- | --- | --- | --- |
|  | tput | median | tput | median | tput | median |
| int8_wo | 0.81× | 0.72× | 0.79× | 0.69× | 0.79× | 0.72× |
| fp8_dyn | 0.50× | 0.30× | 0.71× | 0.44× | **1.01×** | **0.97×** |
| int8_dyn | 0.13× | 0.07× | 0.18× | 0.10× | 0.24× | 0.20× |
| int4_wo | 0.17× | 0.16× | 0.10× | 0.09× | 0.07× | 0.07× |

**Table 14:** Interaction contrasts for int4 wo: the change in effect between scales, from the same protein-clustered paired bootstrap at *R* = 20,000 resamples. The 3B–650M contrast survives Bonferroni correction for all 15 comparisons, where neither individual test does. No other configuration shows a significant interaction, and 15B does not differ significantly from 3B, so the dependence is not monotone in scale.

| Contrast | Difference | 95% CI | $p$ |
| --- | --- | --- | --- |
| 3B – 650M | +0.0101 | [+0.0046, +0.0155] | **0.0007** |
| 15B – 650M | +0.0073 | [+0.0021, +0.0123] | 0.0067 |
| 15B – 3B | −0.0028 | [−0.0062, +0.0006] | 0.1017 |

So the scale dependence of int4 wo is established, while the sign of its effect at any single scale is not: the difference between scales is larger and better resolved than either endpoint. That is the more useful statement anyway, since it is the one that tells a practitioner a result measured at one scale does not transfer.

##### Fidelity predicts magnitude, not direction

Across 3015 assay/configuration pairs, *r*(1 − *ρ*_fp32_, |Δ|) is 0.74, 0.81 and 0.56 at 650M, 3B and 15B. The 0.87 first estimated from 15 pairs sits above all three, which is what a three-assay estimate of a correlation should be expected to do; the bound survives at scale, but weaker than the pilot implied. The *signed* correlation is +0.10, −0.58 and −0.40: it does not hold its sign across scales and is therefore useless for prediction. *ρ*_fp32_ remains the correct cheap screen, bounding how far an answer can move while saying nothing about which way (Figure 3).

**Figure 3:**
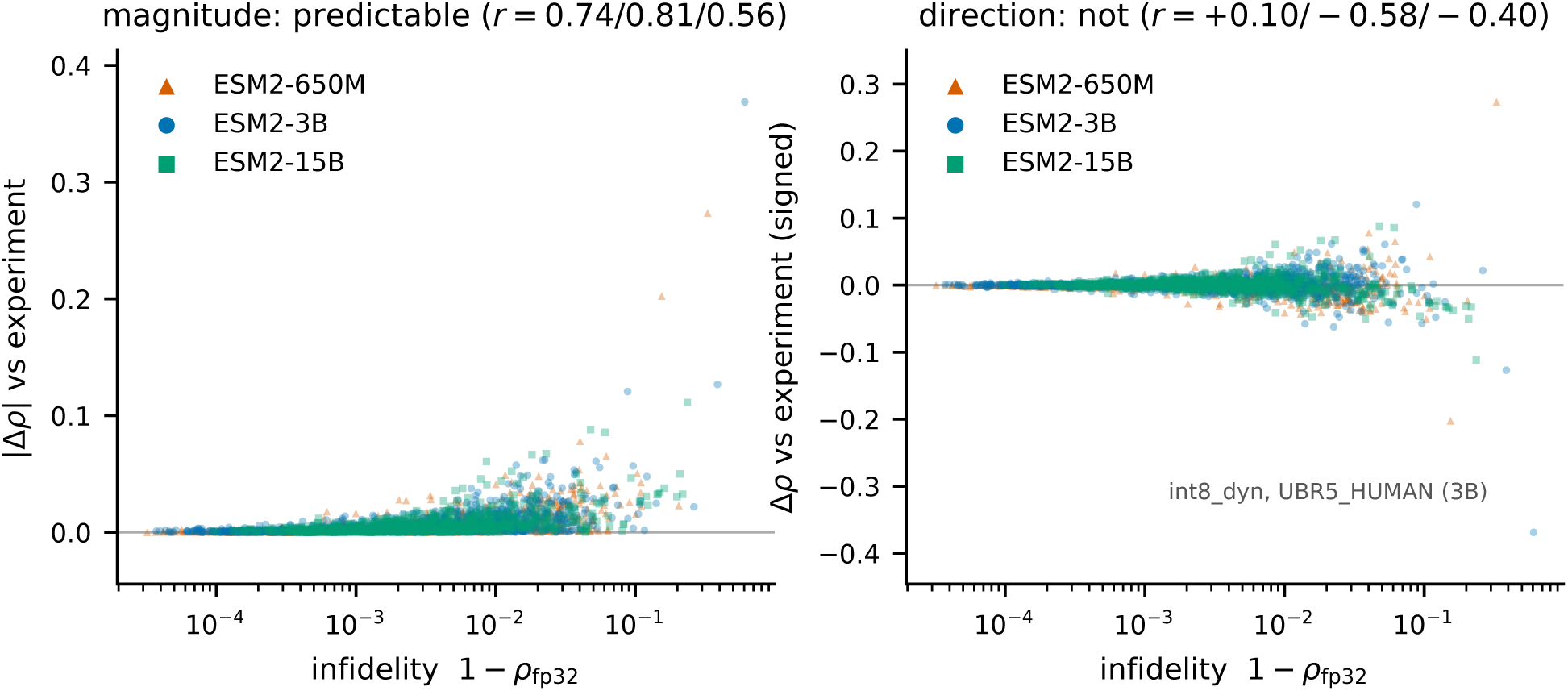
Drift from fp32 against change in ground-truth correlation, over 1005 assay/configuration pairs per model scale. *Left:* infidelity bounds the magnitude of the change — the envelope widens monotonically, and nothing far from fp32 is reliably close to it on the ground truth. *Right:* the same data, signed. The distribution is a symmetric funnel rather than a trend, and the correlation changes sign between scales. A small drift is a valid safety certificate; a large drift is not evidence of a worse model.

**Figure 4:**
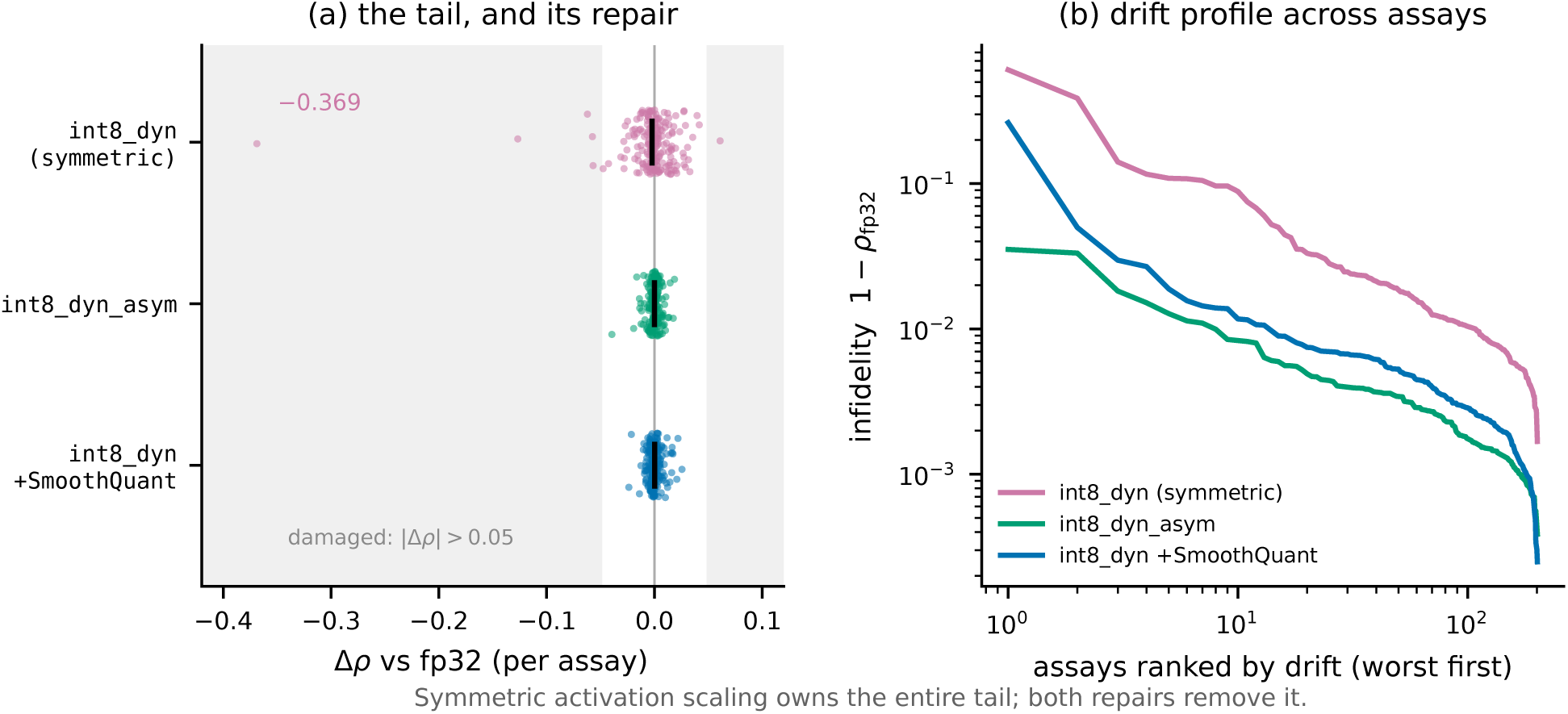
(a) Per-assay change in ground-truth correlation for the three W8A8 variants over all 201 assays at 3B; shaded bands mark |Δ*ρ*| *>* 0.05 and black ticks the benchmark mean. Symmetric activation scaling owns the entire tail, and both repairs remove it while leaving the mean essentially unchanged — the defect is invisible in the statistic most likely to be reported. (b) Drift from fp32 for every assay, ranked worst-first. The separation is not confined to the tail: symmetric scaling is worse across the whole distribution, roughly threefold at the median, which is why its damage is predictable in hindsight and yet absent from its mean.

**Figure 5:**
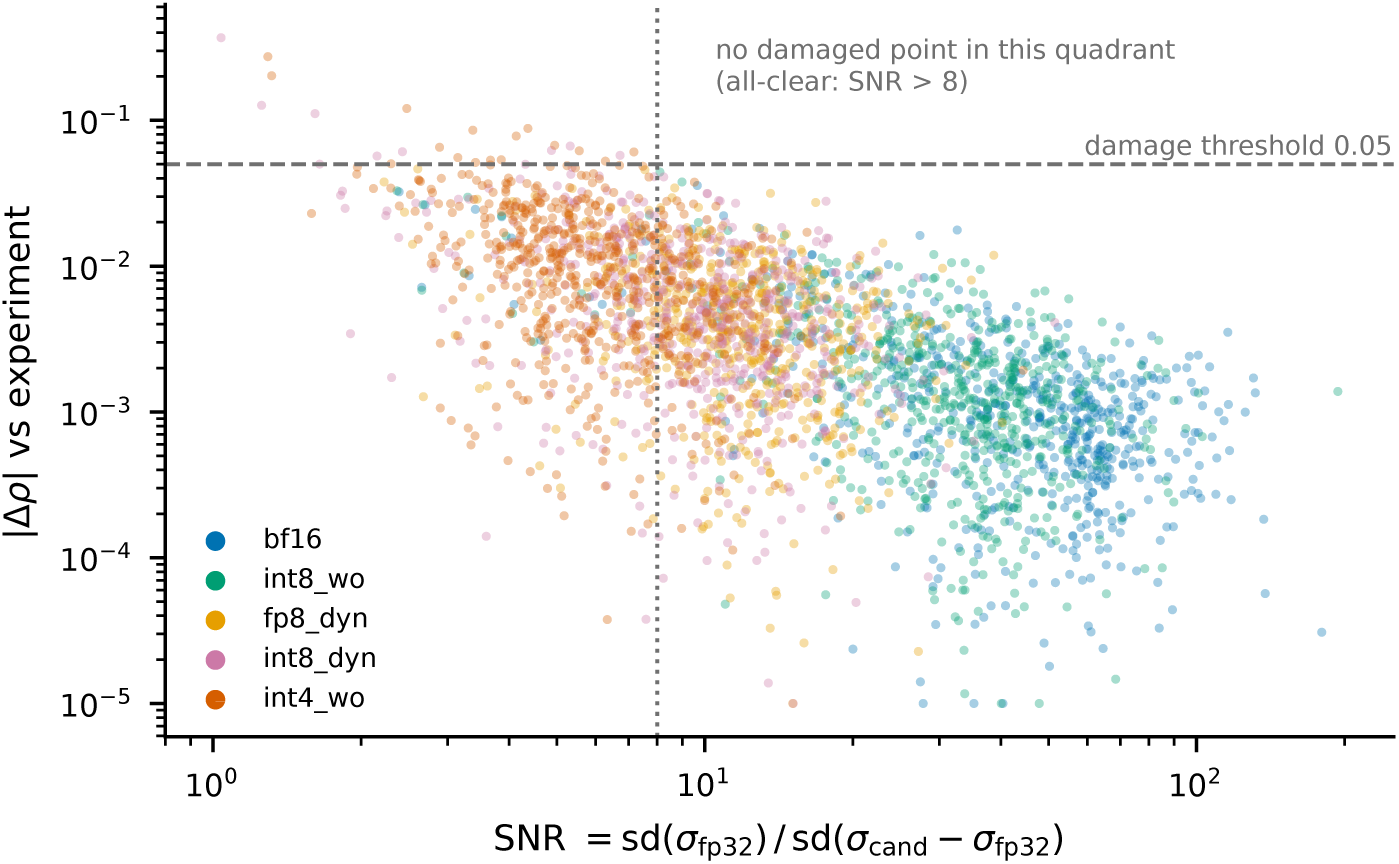
Signal-to-perturbation ratio against realised damage, for all 3015 assay/configuration/scale combinations. The upper-right quadrant — high SNR *and* material damage — is empty: no combination above SNR = 8 loses more than 0.05 Spearman. The relationship is monotone but noisy in the other direction, so a low SNR is a warning rather than a verdict. Colour marks the configuration, and the ordering left-to-right recovers their aggressiveness without being told it, which is a check that the quantity measures what it is supposed to.

##### Stratified effects do not replicate either

At 650M, int4 wo’s cost is monotone in MSA depth — −0.0174, −0.0050, −0.0042 for low, medium and high — which invites the reading that quantization does most harm where the model has least evolutionary signal. That ordering holds at 3B (−0.0011, +0.0007, +0.0097) and breaks at 15B (+0.0011, −0.0009, +0.0033), where every effect is inside the noise band in any case. We report the stratification (Table 15) but draw no conclusion from it: two scales agreeing and a third not is exactly the pattern that the int4 wo result should teach us to distrust.

#### 4.7.6 The W8A8 collapse is a scaling defect, not a limit of the scheme

Reporting that int8 dyn destroys an assay is not the same as establishing that W8A8 cannot work for ESM-2, and the two carry very different recommendations. We tested the difference.

The obvious explanation — too coarse a quantization granularity — is unavailable: int8 dyn already uses *per-token* activation scales and *per-channel* weight scales, so the standard refinement is the configuration already in use. What a per-token scale cannot absorb is a single channel that is large within a token, which is the activation-outlier regime SmoothQuant addresses by migrating outliers into the weights.

We first looked for that signature directly, profiling per-channel activation magnitudes at every encoder Linear input on the collapsing assay and on four length-matched assays that do not collapse. **It is not there.** The collapsing assay’s outlier ratio (median 16.8, p95 310) sits inside the control range (15.5–17.2, 298–457) and its largest raw activation, 14.8, is *smaller* than every control (14.9–17.5). We report this negative result because it is evidence about the diagnostic as much as about the mechanism: aggregating over 180 Linear layers washes out a defect localised to a few, so model-wide activation statistics are not a usable screen.

The intervention test is decisive where the diagnostic was not (Table 16). Both remedies remove every damaged assay.

**Table 15:** int4 wo minus fp32 in mean *ρ*, by ProteinGym function category and by MSA depth, at each scale. The MSA-depth ordering is monotone at 650M and 3B and breaks at 15B; the category ordering does not transfer at all (Binding is the worst stratum at 650M and among the best at 3B). At 15B every effect is inside the noise band of Table 7. We report this for completeness and draw no conclusion from it.

| Stratum | ESM2-650M | ESM2-3B | ESM2-15B |
| --- | --- | --- | --- |
| Activity | −0.0115 | +0.0018 | −0.0013 |
| Binding | −0.0184 | +0.0049 | +0.0004 |
| Expression | +0.0000 | −0.0018 | +0.0015 |
| OrganismalFitness | −0.0117 | −0.0003 | −0.0001 |
| Stability | +0.0025 | +0.0101 | +0.0032 |
| MSA depth: low | −0.0174 | −0.0011 | +0.0011 |
| MSA depth: medium | −0.0050 | +0.0007 | −0.0009 |
| MSA depth: high | −0.0042 | +0.0097 | +0.0033 |

**Table 16:**
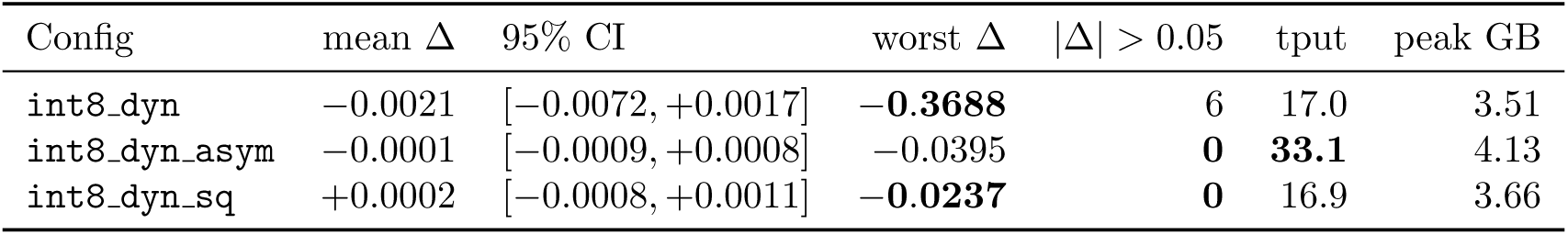
Repairing W8A8 at 3B over the full benchmark. int8 dyn asym is the same configuration with asymmetric rather than symmetric activation quantization — one keyword argument, no calibration. int8 dyn sq adds SmoothQuant with *α* = 0.5 calibrated on 16 target sequences. Both eliminate all six damaged assays and cut the worst case by an order of magnitude. The defect is therefore in the default activation scaling, not in W8A8. Intervals are the same protein-clustered paired bootstrap used throughout (*R* = 20,000); none of the three differs significantly from fp32 on the *mean*, which is exactly the point — the mean cannot separate them and the worst case can. Worst-assay fidelity *ρ*_fp32_ is 0.393, 0.965 and 0.737 respectively; *tput* is aggregate masked positions per second, i.e. total positions over total seconds. These two remedy comparisons bring the family of tests against fp32 to 17, giving a Bonferroni threshold of 0.0029; the conclusions in Section 4.7 are unchanged by the revised denominator.

**Table 17:** SNR against realised damage, over all 3015 assay/configuration/scale combinations. Every collapse beyond 0.10 in this study has SNR *<* 1.7, and no combination with SNR *>* 8 is damaged beyond 0.05. The 0.0% is 0 of 2233, which bounds the underlying rate at 0.13% with 95% confidence rather than establishing it as zero. The screen is therefore usable as an all-clear rather than as a precise predictor: a high SNR licenses deployment, a low one demands a closer look.

| SNR | $n$ | median $\vert \Delta\vert $ | $P(\vert \Delta\vert > 0.05)$ | worst $\Delta$ |
| --- | --- | --- | --- | --- |
| $< 2$ | 13 | 0.0478 | 46.2% | -0.3688 |
| 2-4 | 139 | 0.0165 | 7.2% | -0.0569 |
| 4-8 | 630 | 0.0092 | 1.4% | -0.0622 |
| $> 8$ | 2233 | 0.0019 | **0.0%** | -0.0409 |

On the two assays that collapse, int8 dyn’s −0.3688 and −0.1268 become +0.0102 and −0.0164 under asymmetric scaling and +0.0034 and −0.0098 under SmoothQuant, with fidelity against fp32 on the worst assay rising from 0.393 to 0.999.

The practical ordering is not the one we expected. **Asymmetric activation quantization is the better remedy**: it needs no calibration data, no extra pass and no prototype dependency, and in our measurements it ran 1.95× faster than symmetric int8 dyn (20.6 against 40.0 minutes for the benchmark) at 0.6 GB more peak memory. We did not investigate the speed difference and do not claim it generalises. SmoothQuant achieves a slightly better worst case (−0.024 against −0.040) but costs a calibration pass, and its worst-assay *fidelity* is actually poorer (0.737 against 0.965).

This changes a recommendation. “Avoid int8 dyn” was too general: the correct statement is that its default *symmetric* activation scaling is unsafe on ESM-2 and that a one-line change repairs it. Symmetric scaling allocates half the INT8 range to values of a sign that post-GELU activations rarely take, which is a plausible account of why the asymmetric variant helps, though we have not isolated it.

#### 4.7.7 The repair helps accuracy on both workloads and speed on only one

The asymmetric variant was adopted above on DMS evidence, where it is both more accurate and 1.95× faster than symmetric. Applying that conclusion to bulk embedding without measuring it would repeat the error this paper is about, so we measured it at the settings of Table 5, where it appears as its own row.

The *accuracy* repair transfers. Embedding cosine rises from 0.9896 under symmetric scaling to 0.9995, logit KL falls from 1.4 × 10^−2^ to 1.4 × 10^−3^, and DMS *ρ*_fp32_ from 0.9928 to 0.9972. Whatever symmetric scaling is doing wrong, it is not specific to the masked-marginals input distribution.

The *speed* effect reverses. Asymmetric runs at 0.90× bf16 against symmetric’s 1.05× — a 14% throughput loss — having been 1.95× *faster* than symmetric on DMS scoring. The same one-keyword change is a large speed win on one workload and a moderate loss on the other, which is the workload-dependence of this paper’s central claim reappearing inside a single configuration.

The practical consequence corrects a recommendation we made from the DMS result alone. For bulk extraction on hardware without FP8, int8 wo **beats the asymmetric variant on both axes that matter there**: faster (0.96× against 0.90×) and more faithful (embedding cosine 0.999956 against 0.999469), at 1.22 GB more peak memory. Asymmetric int8 dyn is worth its throughput cost only where the memory is needed and FP8 is unavailable.

We were unable to obtain the corresponding A100 measurement — that partition was occupied by a multi-day job for the duration of this revision — so the ordering above is established on H200 and the A100-specific numbers remain unmeasured. The recommendation for FP8-less hardware is stated as an inference from this ordering rather than as a measurement, and marked as such in Table 23.

#### 4.7.8 A label-free screen for which assays are at risk

Chasing this mechanism produced something more useful than the mechanism. What determines whether a ranking survives a perturbation is not the size of the perturbation but its size *relative to the spread of the scores being perturbed*. Both quantities are computable from the candidate and the fp32 reference on a user’s own target, with no experimental labels:

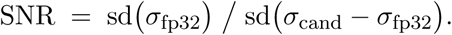

The obvious objection is that SNR simply re-discovers the configuration ranking — median SNR runs from 47.9 for bf16 down to 5.7 for int4 wo — in which case the bins are sorting configurations, not targets, and the screen says nothing that choosing bf16 would not. It does not. Holding the configuration fixed, SNR still predicts damage within it (Table 18), and within int4 wo alone — the configuration with the broadest tail — the probability of losing more than 0.05 Spearman falls from 10.5% below SNR = 4 to 1.6% between 4 and 8 and to 0.0% above 8 (95, 367 and 141 assay/scale combinations respectively). The quantity therefore carries per-target information, which is what a screen has to do to be worth running.

**Table 18:** SNR against damage *within* each configuration. If the screen merely recovered the aggressiveness ordering these correlations would be near zero; they are not. The negative sign is the expected direction: more signal relative to perturbation, less damage.

| Config | median SNR | Spearman(SNR, $\vert \Delta\rho\vert $) within config |
| --- | --- | --- |
| bf16 | 47.9 | −0.425 |
| int8_wo | 34.6 | −0.404 |
| fp8_dyn | 11.9 | −0.264 |
| int8_dyn | 10.1 | −0.386 |
| int4_wo | 5.7 | −0.389 |

**Table 19:**
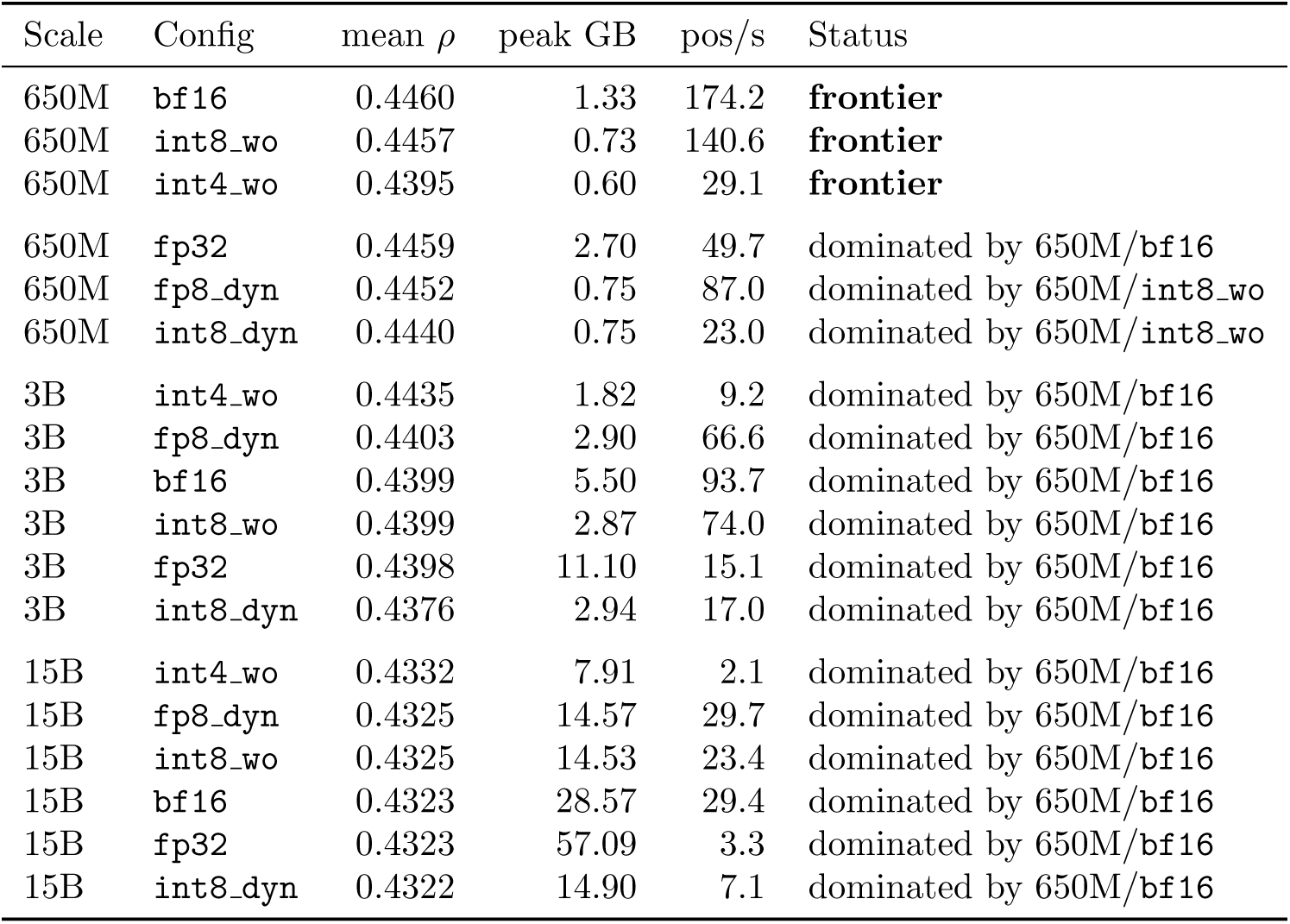
Cross-scale Pareto surface, all eighteen scale/configuration combinations, ordered by accuracy. An option is dominated if another is at least as good on accuracy, peak memory *and* speed, and strictly better on one; the dominator named is in every case itself on the frontier. Speed is aggregate throughput — total masked positions over total seconds — and the frontier is unchanged if the median-of-assay convention is substituted — the same three options are non-dominated either way, so this result does not rest on that choice. Every 3B and 15B configuration is dominated by a 650M one, including 3B/int4 wo, which our own recommendations table previously offered for minimum memory and which uses *more* memory than 650M/bf16 while being less accurate and 19× slower.

| Scale | Config | mean $\rho$ | peak GB | pos/s | Status |
| --- | --- | --- | --- | --- | --- |
| 650M | `bf16` | 0.4460 | 1.33 | 174.2 | **frontier** |
| 650M | `int8_wo` | 0.4457 | 0.73 | 140.6 | **frontier** |
| 650M | `int4_wo` | 0.4395 | 0.60 | 29.1 | **frontier** |
| 650M | `fp32` | 0.4459 | 2.70 | 49.7 | dominated by 650M/`bf16` |
| 650M | `fp8_dyn` | 0.4452 | 0.75 | 87.0 | dominated by 650M/`int8_wo` |
| 650M | `int8_dyn` | 0.4440 | 0.75 | 23.0 | dominated by 650M/`int8_wo` |
| 3B | `int4_wo` | 0.4435 | 1.82 | 9.2 | dominated by 650M/`bf16` |
| 3B | `fp8_dyn` | 0.4403 | 2.90 | 66.6 | dominated by 650M/`bf16` |
| 3B | `bf16` | 0.4399 | 5.50 | 93.7 | dominated by 650M/`bf16` |
| 3B | `int8_wo` | 0.4399 | 2.87 | 74.0 | dominated by 650M/`bf16` |
| 3B | `fp32` | 0.4398 | 11.10 | 15.1 | dominated by 650M/`bf16` |
| 3B | `int8_dyn` | 0.4376 | 2.94 | 17.0 | dominated by 650M/`bf16` |
| 15B | `int4_wo` | 0.4332 | 7.91 | 2.1 | dominated by 650M/`bf16` |
| 15B | `fp8_dyn` | 0.4325 | 14.57 | 29.7 | dominated by 650M/`bf16` |
| 15B | `int8_wo` | 0.4325 | 14.53 | 23.4 | dominated by 650M/`bf16` |
| 15B | `bf16` | 0.4323 | 28.57 | 29.4 | dominated by 650M/`bf16` |
| 15B | `fp32` | 0.4323 | 57.09 | 3.3 | dominated by 650M/`bf16` |
| 15B | `int8_dyn` | 0.4322 | 14.90 | 7.1 | dominated by 650M/`bf16` |

**Table 20:**
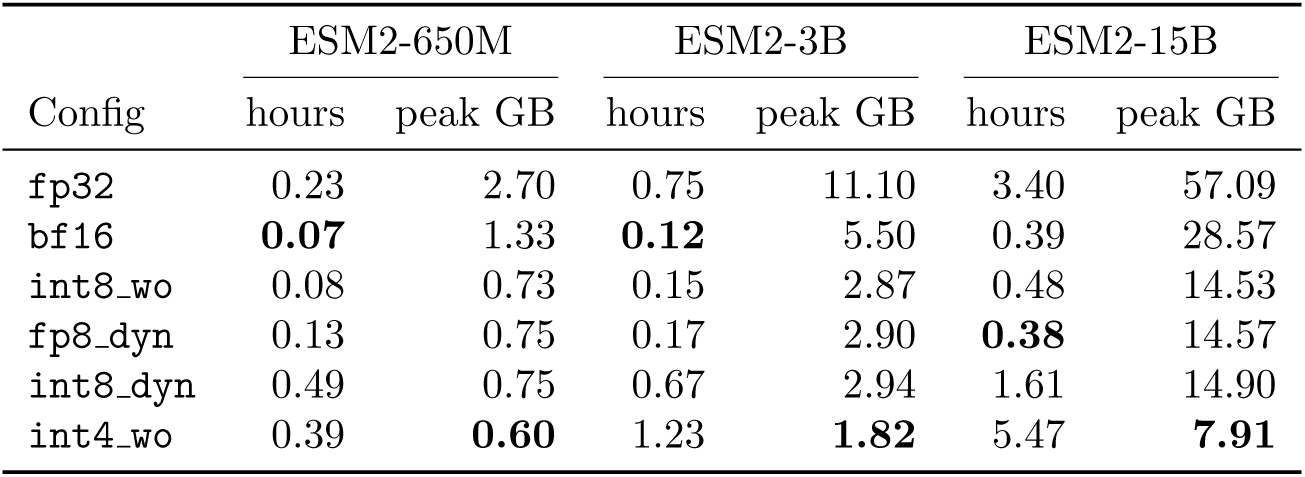
Full 201-assay benchmark by model scale, uncompiled, one H200. Wall-clock is summed scoring time — the quantity the throughput ratios of Table 13 are consistent with — and peak memory is the median across assays. fp32 at 15B needs 57 GB, so the reference configuration alone rules out an 80 GB A100 once activations are included.

| Config | ESM2-650M |  | ESM2-3B |  | ESM2-15B |  |
| --- | --- | --- | --- | --- | --- | --- |
|  | hours | peak GB | hours | peak GB | hours | peak GB |
| fp32 | 0.23 | 2.70 | 0.75 | 11.10 | 3.40 | 57.09 |
| bf16 | **0.07** | 1.33 | **0.12** | 5.50 | 0.39 | 28.57 |
| int8_wo | 0.08 | 0.73 | 0.15 | 2.87 | 0.48 | 14.53 |
| fp8_dyn | 0.13 | 0.75 | 0.17 | 2.90 | **0.38** | 14.57 |
| int8_dyn | 0.49 | 0.75 | 0.67 | 2.94 | 1.61 | 14.90 |
| int4_wo | 0.39 | **0.60** | 1.23 | **1.82** | 5.47 | **7.91** |

**Table 21:** Compilation cost inside the timed region. A single forward pass at the DMS input shape costs 15 s to autotune and 28 ms to execute, so a timed region containing compilation ranks configurations by their number of Triton kernel candidates rather than by their speed.

| Measurement | Time |
| --- | --- |
| eager, one forward at shape (11, 42) | 27.9 ms |
| eager, full `masked marginals` pass | 0.03 s |
| compiled, one forward at shape (11, 42) | **15,138.1 ms** |

**Table 22:** The peak-memory artifact. Each configurations measured peak contains the previous configurations weights, which surfaces as bf16 apparently needing more peak memory than fp32.

| Config | Weights (GB) | Peak (GB) | Implied overhead |
| --- | --- | --- | --- |
| fp32 | 2.49 | 2.56 | +0.07 |
| bf16 | 1.21 | 3.74 | +2.53 ≈ fp32’s weights |
| int8_wo | 0.61 | 1.86 | +1.25 ≈ bf16’s weights |
| int8_dyn | 0.61 | 1.26 | +0.65 ≈ int8_wo’s weights |

**Table 23:** Recommendations by workload and binding constraint. Model scale is the first decision rather than a column, because the cross-scale Pareto surface (Table 19) shows every 3B and 15B configuration to be dominated by a 650M one for this workload. Two rows were wrong in earlier drafts and are corrected here on measured evidence: 3B/int4 wo for minimum memory (it uses more memory than 650M/bf16, is less accurate and is 19× slower), and asymmetric int8 dyn for FP8-less bulk extraction (it is slower *and* less faithful than int8 wo there, Table 5).

| Workload | Constraint | Recommendation |
| --- | --- | --- |
| Variant-effect scoring | accuracy | `bf16` at **650M** — 3B and 15B are not better and cost more |
| Variant-effect scoring | memory | 650M/`int8_wo` (0.73 GB) before any quantization of a larger model |
| Variant-effect scoring | 3B/15B fixed | `bf16` up to 3B; `fp8_dyn` at 15B |
| Bulk embedding (H200) | throughput | `fp8_dyn` + `max-autotune-no-cudagraphs` |
| Bulk embedding (A100) | throughput | `int8_wo`<sup>†</sup> — faster *and* more faithful than asymmetric `int8_dyn` on this workload; avoid symmetric |
| Minimum footprint | memory | 650M/`int4_wo` (0.60 GB), only after checking drift on *your* target: worst −0.20, or −0.05 on assays with signal |
| QLoRA fine-tuning base | memory | `int4_wo` at the scale being tuned |
| V100 / sm_70 | any | Quantization buys memory only, never speed |

<sup>†</sup> Inferred from the H200 ordering in Table 5. The A100 partition was occupied by a multi-day job for the duration of this revision, so the A100-specific bulk numbers are unmeasured.

This is a better screen than the raw drift test in Algorithm 3, because it is calibrated: *ρ*_fp32_ bounds how far an answer can move without saying whether that distance matters for the assay at hand, whereas SNR compares the movement to the signal it has to preserve. We would now run both.

#### 4.7.9 Quantization competes with using a smaller model, and loses

Every comparison so far is within a scale: a configuration against fp32 on the same checkpoint. That is the wrong frame for a practitioner, who is not obliged to run 3B at all. Once the three scales are placed on one Pareto surface over (accuracy, peak memory, speed), **only three of the eighteen configurations are non-dominated, and all three are 650M** (Figure 6, Table 19).

**Figure 6:**
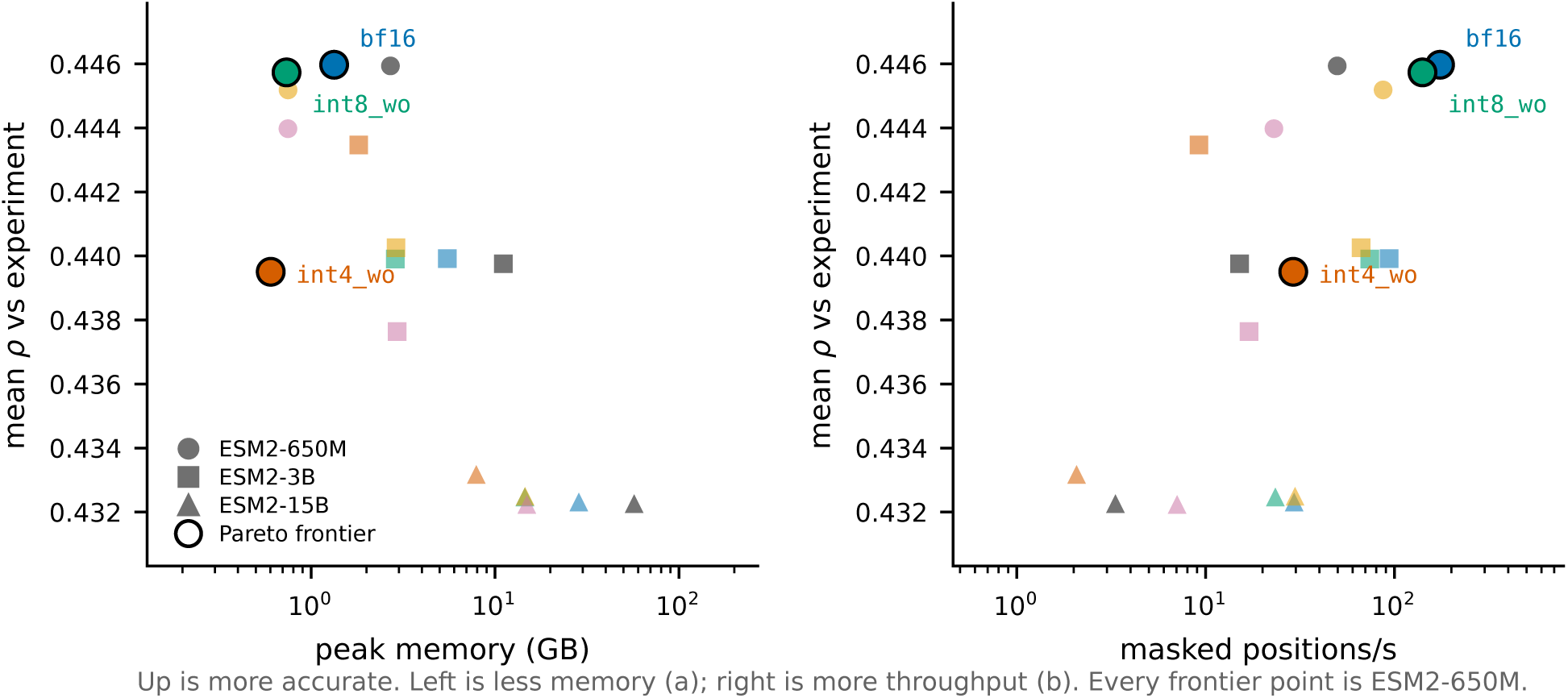
All eighteen scale/configuration combinations on two projections of the Pareto surface. Higher is more accurate in both panels; the horizontal axes run in opposite senses, so better is leftward against peak memory (a) and rightward against throughput (b). Circled points are non-dominated, and all are ESM2-650M. The practical consequence is that aggressive quantization of a large model is not a way to buy a small memory footprint for this workload — a small model already provides one, more accurately and far faster.

We take this to be the most consequential practical result in the paper, and it is invisible to a within-scale comparison. Quantization remains useful *within* the frontier model: 650M/int8 wo halves the memory of 650M/bf16 for a 0.0002 accuracy change, and 650M/int4 wo halves it again.

But the memory-oriented configurations at 3B and 15B are not trade-offs against a small model; they are strictly worse than one.

Two qualifications. First, this holds for variant-effect scoring, where 650M is statistically indistinguishable from 3B and 15B on ground truth (Section 4.7); it does not transfer to tasks where the larger checkpoints are genuinely better, and structure prediction is the obvious such task. Second, dominance is stated on *benchmark-mean* accuracy, and Section 4.7.1 argues that means are the wrong selection criterion — a practitioner with a specific target should check the tail rather than read the frontier off this table.

#### 4.7.10 Speed and memory over the whole benchmark

The DMS recommendation from three assays survives at 650M and 3B and inverts at 15B. Up to 3B, bf16 eager is fastest at every problem size — 6.2× faster than fp32 end-to-end at 3B — with ground-truth agreement never moving more than 0.018 across 402 assay/model pairs, and int8 wo is the memory fallback at roughly 80% of bf16 throughput. At 15B, fp8 dyn matches bf16 on speed at half the memory and becomes the default. int8 dyn and int4 wo are not trade-offs for this workload at any scale: they are slower than bf16 *and* carry the two worst tails in Table 8.

## 5 Reproducibility Pitfalls in Quantization Benchmarking

We document four measurement defects encountered during this study. They are included in the main text, unusually for a paper, because they bear directly on its thesis: three of the four produced clean, monotonic, plausible result tables while measuring the wrong quantity, and each would have supported a confident and wrong recommendation. A reader who accepts our argument that benchmark averages can mislead should also want to know how the underlying measurements can mislead. All four are properties of the standard tooling rather than of this codebase, and we expect them to recur in any comparable study.

### 5.1 Compile cost inside the timed region (three occurrences)

The DMS throughput column initially ranked fp32 fastest and fp8 dyn slowest — an exact inversion of the bulk-throughput column. Two successive fixes failed. Direct instrumentation finally established the cause. On 650M/A100 with a 40-residue wild type over 11 masked positions:

DMS scoring uses a much smaller input shape than the bulk-throughput batches, and under max-autotune each new shape costs roughly 15 s to autotune. At this batch size that cost recurs rather than amortizing. The timed region was therefore measuring *autotune cost*, and ranking configurations by their number of Triton kernel candidates — fp32 had the fewest and appeared fastest, fp8 dyn had 98 and appeared slowest. The fix was to score DMS against the uncompiled module.

This has a consequence beyond the benchmark: for short wild-type sequences with few mutated positions, max-autotune is actively counterproductive in production, not merely in measurement.

The same class of defect appeared earlier in the bulk-throughput path, where warming up on only a prefix of the batches left recompilation for unseen length buckets inside the timed region.

### 5.2 Peak memory attributed to the wrong model

Peak memory during DMS scoring was reported as follows, with the activation overhead implied by subtracting weights:

Each configuration’s peak included the *previous* configuration’s weights, producing the visible absurdity of bf16 requiring more peak memory than fp32. The cleanup helper deleted only its own local reference to the model; the caller retained a binding. Because nn.Module graphs contain reference cycles, dropping the last reference does not free them by reference counting alone — they persist as cyclic garbage until a collection runs. Inserting an explicit gc.collect() before each peak-memory reset corrected it; overhead became a consistent 0.04–0.07 GB.

The bulk-throughput memory column was verified unaffected: bf16’s measured overhead there was +7.34 GB, which cannot contain fp32’s 10.72 GB of weights.

### 5.3 Silent argument leakage in the job script

A batch script used shift 3 || true. When fewer than three positional arguments are supplied, shift 3 || fails, true suppresses the failure, and the arguments remain in “$@” — where they were appended to the Python invocation as stray arguments. Relying on a default for the third argument was sufficient to trigger it.

### 5.4 Common thread

Three of the four defects shared a signature: a cost that belonged outside the measurement leaked inside it, and the resulting ranking was plausible enough to be believed. In each case the tell was a result that inverted or contradicted a neighbouring measurement. We would emphasise that a benchmark producing a *clean, monotonic, plausible* table is not thereby correct; the memory bug produced exactly such a table for several runs.

## 6 Discussion

### 6.1 The two workloads want opposite configurations

This is the principal practical finding.

**Bulk embedding extraction** is compute-bound: large batches, large GEMMs. Quantization pays. fp8 dyn with max-autotune wins on all three axes — 1.15× bf16 throughput, weights 5.29 → 2.65 GB, peak 12.63 → 7.24 GB, with accuracy indistinguishable from bf16.

**DMS variant-effect scoring** is latency-bound: the masked batch is small, no GEMM is large enough for low precision to help, and per-linear quantize/dequantize overhead is pure loss. bf16 in eager mode is fastest at every problem size, and max-autotune is actively harmful.

Applying the embedding-optimized configuration to DMS scoring makes it up to an order of magnitude slower. Because both workloads commonly appear in the same pipeline, this is a live risk rather than a theoretical one. Figure 7 shows the two frontiers side by side.

**Figure 7:**
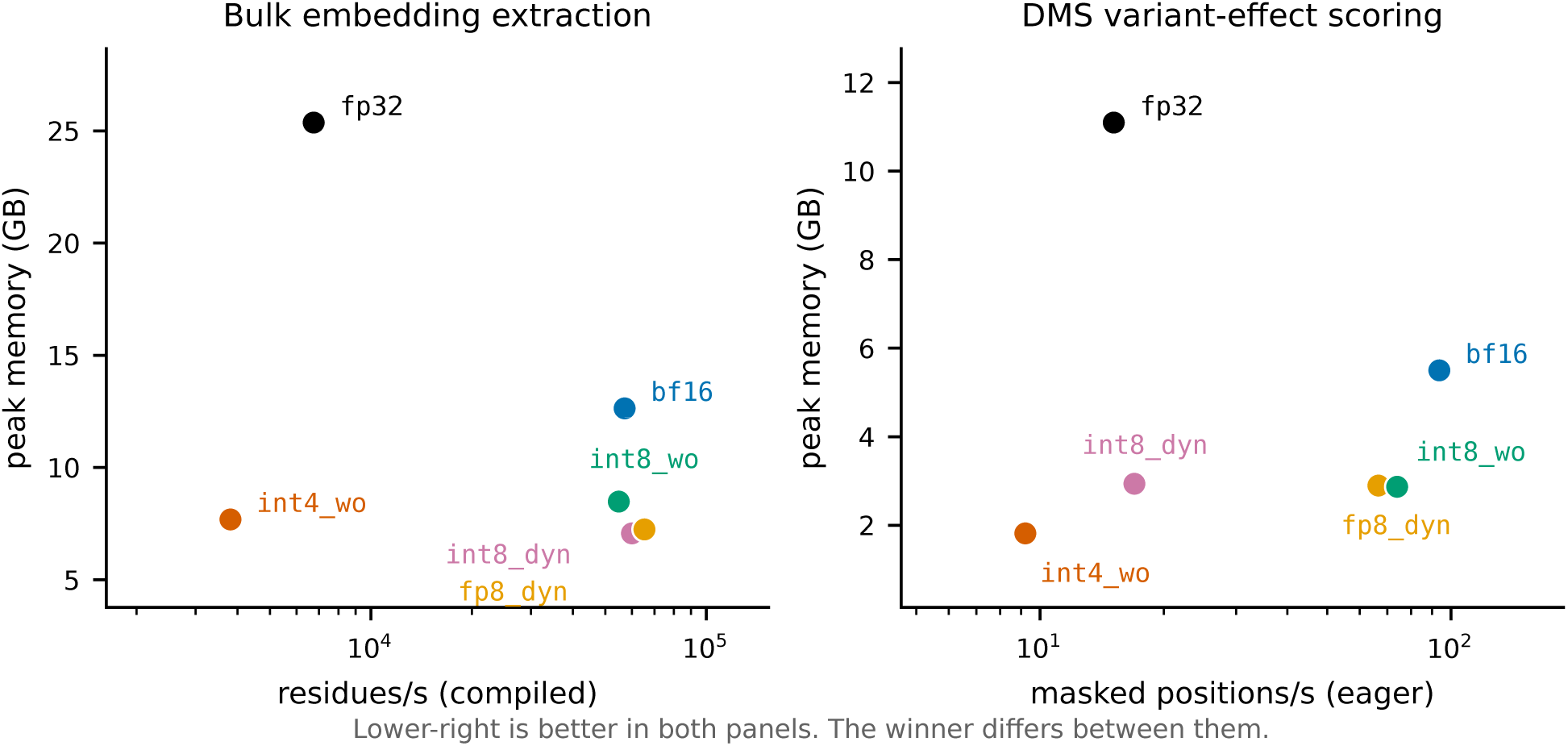
Speed–memory frontiers for the two workloads, ESM2-3B on one H200; lower-right is better in both panels. *Left:* bulk embedding extraction under max-autotune, where fp8 dyn is fastest and near-smallest. *Right:* DMS scoring over the full 201-assay benchmark in eager mode, where the same configuration is 1.4× slower than bf16 and the ranking is substantially reordered. No single configuration is preferred by both.

**This conflict is bounded by scale, and we would not have known that from one model** (The evidence is one-sided: the DMS side of the comparison spans three scales, the bulk-embedding side is measured at 3B only.) The mechanism is that fp8 dyn’s per-call quantization overhead is fixed while the useful work grows with hidden size, so the penalty shrinks as the model grows: 0.50× bf16 DMS throughput at 650M, 0.71× at 3B, 1.01× at 15B (Table 13; the ordering is the same under the median-of-assay convention, at 0.30, 0.44 and 0.97). By 15B the DMS batch is large enough in the hidden dimension to be compute-bound despite being narrow in the batch dimension, and fp8 dyn becomes the right choice for *both* workloads at half the memory. The general lesson is not that DMS is latency-bound — it is that whether a workload is latency-bound is a property of the model and the workload together, not of the workload alone.

### 6.2 On the interpretation of fidelity metrics

The conventional validation practice — quantize, measure drift against the full-precision model, accept if drift is small — is sound as a *risk bound*. Our data support the bound at scale: over 3015 assay/configuration pairs, fidelity loss correlates with the magnitude of ground-truth change at *r* = 0.74, 0.81 and 0.56 at 650M, 3B and 15B — the same relationship the 15-pair pilot found at *r* = 0.87, weaker at every scale once the sample is large enough to resolve it.

It is not sound as a *quality ranking*. A configuration with larger drift is not thereby less accurate against reality; it is merely less predictable. The signed correlation is +0.10, −0.58 and −0.40 at 650M, 3B and 15B — it does not hold its sign across model scales.

A tempting misreading of our BLAT result is that quantization acts as beneficial regularization. We do not endorse this, and the full benchmark settles it: int4 wo is significantly *negative* at 650M, significantly *positive* at 3B, and null at 15B. Two scales would have permitted the reading that sensitivity falls with model size; the third removes it. A configuration cannot be genuinely more accurate than the reference from which it is derived in any way one could rely on in advance.

### 6.3 A failure worth diagnosing is worth trying to fix

The first version of this study stopped at “int8 dyn destroys one assay, so avoid it”. That is a defensible reading of the benchmark and it is wrong in a way that matters: the collapse is a property of one scaling choice inside the configuration, not of the configuration, and certainly not of W8A8 on encoders. Switching the activation mapping from symmetric to asymmetric — a keyword argument — removes every damaged assay (Section 4.7.6).

We think the general lesson is about what a benchmark result licenses. Measuring that a configuration fails establishes that it fails *as configured* ; it does not establish that the approach is unusable, and the distance between those two claims is one experiment. A study that reports only the ranking leaves the reader unable to tell which of them it has shown. It is worth noting how close we came to publishing the stronger claim: the collapse is dramatic, the explanation “W8A8 is too aggressive for this model” is available and plausible, and nothing in the benchmark contradicts it.

The diagnostic that failed is instructive too. We looked for the textbook activation-outlier signature on the collapsing assay and did not find one — its largest activation is *smaller* than every control’s. Had we stopped there we would have concluded that outliers were not the mechanism, and the successful repair by two methods that both target activation scaling shows that conclusion would also have been wrong. The diagnostic was averaging over 180 Linear layers, which is the wrong resolution for a defect concentrated in a few. Negative results from aggregate statistics are weak evidence about localised phenomena, and we report ours as such rather than as a finding.

### 6.4 Averages conceal the failure that matters

The benchmark-mean result is that no configuration changes mean ground-truth correlation by more than 0.007. Read alone, that licenses any configuration on the list. It should not.

int8 dyn at 3B is statistically indistinguishable from fp32 on the mean (*p* = 0.34) and takes one assay from *ρ* = 0.591 to 0.223. A practitioner does not deploy against the mean of 201 assays; they deploy against one protein. The benchmark mean answers whether a configuration is safe *on average over targets*, which is a claim about a population, while the operative question is whether it is safe on a specific target.

The practical consequence is that the tail statistics — worst-case single-assay drift, and the count of assays exceeding a tolerance — are the numbers to select on, and they separate the configurations far more sharply than the means do. On that criterion bf16 and int8 wo are safe (no assay beyond 0.018 across 402 assay/model pairs), fp8 dyn is nearly so (0.034), and int8 dyn and int4 wo are not.

### 6.5 Does the repair undercut the argument?

An objection follows directly from Section 4.7.6, and we think it is the strongest one available against this paper. The failure that motivates the whole mean-versus-tail argument turns out to be repairable by a keyword argument. So did the tail analysis discover a property of quantization, or merely a configuration bug that a careful practitioner would have found by trying the other activation mapping? On that reading the title promises something the paper does not deliver: what benchmark averages concealed was an implementation defect, not a failure that matters.

Three things are worth separating in reply.

First, **the repair is only reachable through the tail**. Nobody switches an activation mapping that shows no symptom, and int8 dyn shows none in the statistic that would normally be reported: its benchmark mean is −0.0021 with *p* = 0.34, indistinguishable from fp32, and it stays indistinguishable after the repair. A study reporting means would have shipped the broken configuration as safe, and would have had no reason to look for a fix. That the defect turned out to be cheap to correct once seen is an argument for looking, not against it.

Second, **not every tail is a bug**. int4 wo damages 4, 5 and 2 of the ∼157 signal-carrying assays by more than 0.05 at 650M, 3B and 15B respectively, with worst cases of −0.050, −0.056 and −0.047 — a tail present at every scale we measured. We know of no analogous one-line repair for it, and its damage is distributed across assays and scales rather than concentrated in a single collapse. The two configurations therefore fail in different ways — one sharply and repairably, one broadly and persistently — and the benchmark mean is equally blind to both. Had int8 dyn been the only case, the objection would carry more weight than it does.

Third, and conceding the substance of it: **the mean cannot tell you which kind of failure you have**. It reports that a configuration is safe on average over targets; it cannot distinguish a repairable defect from an intrinsic limit, because it does not surface either. The honest form of our claim is therefore narrower than the title alone suggests. Benchmark averages conceal *heterogeneity*, and whether a particular instance of heterogeneity is a bug worth fixing or a property worth avoiding is a second question, which the tail statistics raise and do not answer. We would rather state the claim at that strength than defend the stronger one.

### 6.6 Recommendations

Two non-quantization measures outweigh most of the above. Moving from fp32 to bf16 is worth 8.4× on bulk extraction and 6.2× over the full DMS benchmark; length-bucketed batching eliminates 25% of wasted computation. Both should be applied before any quantization is considered.

A third, proposed in an earlier draft of this report, does not survive the full benchmark: the 650M checkpoint beat 3B by 0.14 Spearman on BLAT, which suggested model selection dominated precision selection. Over 201 assays the two are indistinguishable (0.4459 vs 0.4398, 95% CI [−0.0061, +0.0185], *p* = 0.35, with 650M ahead on 110 of 201). What remains true is narrower and still useful: the *uncertainty* on model choice (±0.012) exceeds every quantization effect measured here (≤ 0.007). Precision is the settled variable; checkpoint choice is not, and is worth benchmarking on a representative assay.

### 6.7 A selection protocol

Our results imply a procedure rather than a ranking, because the quantity that separates configu-rations — worst-case single-assay drift — is cheap to measure on a target and cannot be inferred from published means. Algorithm 3 states it. The essential point is line 4: the screen runs against the fp32 model on the user’s own sequences and needs no experimental labels, so it costs one extra scoring pass and no wet-lab work.

**Algorithm 3** Choosing a precision configuration for a variant-effect workload. The screen requires no ground-truth labels, because Section 6.2 establishes that drift from fp32 bounds how far the answer can move even though it does not indicate direction. The SNR test added at the third line is what converts that bound into a decision: drift alone is uncalibrated, since the same absolute movement is harmless on an assay with a wide score distribution and fatal on a narrow one (Section 4.7.8).

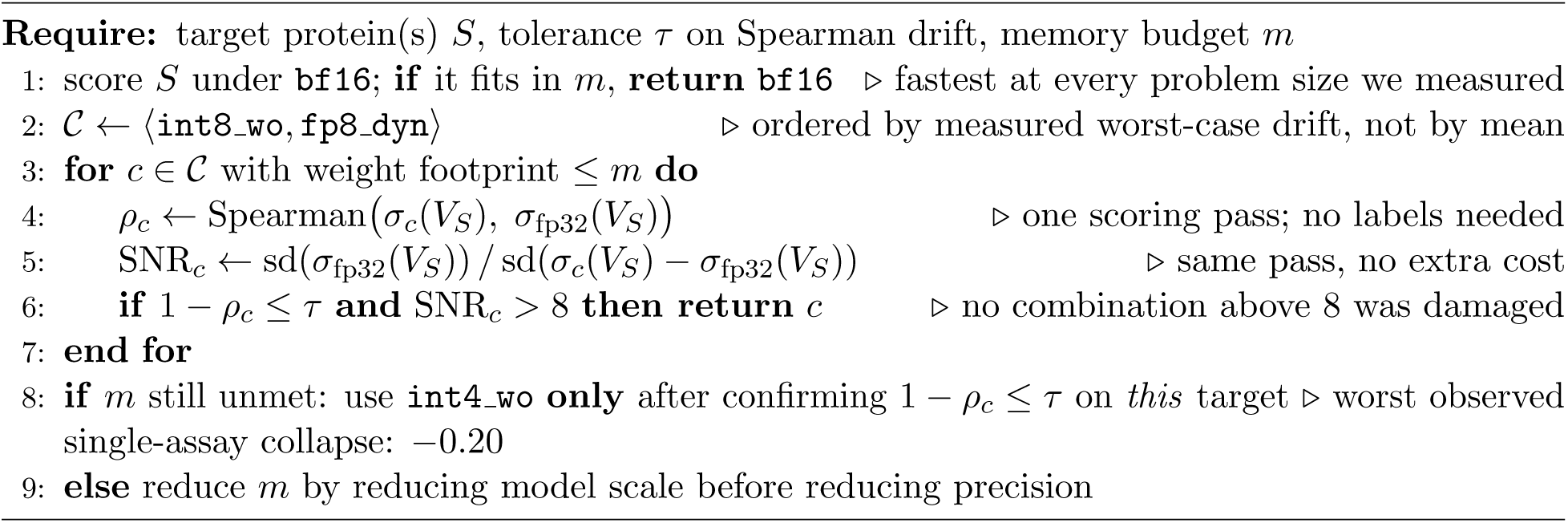

[uWe deliberately do not recommend selecting on published benchmark means. On the 201-assay mean, int8 dyn and bf16 are separated by 0.002; on worst-case single-assay behaviour they are separated by 0.36. The mean is the statistic most likely to be quoted and the least useful for this decision.

### 6.8 Computational cost

The full benchmark costs 1.4 GPU-hours at 650M, 3.1 at 3B and 11.7 at 15B on a single H200 — 40,775 masked forward passes per configuration, six configurations, three scales, 16.2 GPU-hours in total. Cost is dominated by the two configurations we recommend against: int4 wo alone is 5.5 of the 11.7 hours at 15B. Restricting to fp32 plus the three viable configurations would cost 4.7 hours at 15B and 6.4 overall.

This is small enough that evaluating on the complete benchmark rather than a subset is a matter of choosing to, not of affording to. The three-assay evaluation this study began with saved roughly two GPU-hours and produced two conclusions that the full benchmark reversed; running one scale rather than three would have saved a further thirteen and left a third.

## 7 Limitations

- **We still cannot predict which assays behave which way.** 201 assays establish the population behaviour of each configuration and the size of its tail. They do not identify in advance which target will be the one that collapses. The only reliable procedure remains scoring a candidate configuration against fp32 on the actual target.
- **Three scales are enough to refute a trend, not to establish one.** We can say the int4 wo effect has no monotone dependence on model size, and that the workload conflict dissolves somewhere between 3B and 15B. We cannot say where that boundary lies, nor whether the fp8 dyn crossover continues past 15B, because ESM-2 has no larger checkpoint.
- **15B is less numerically stable even at bf16.** Its worst-case bf16 drift is −0.0323 against −0.0072 at 3B (Table 8). We did not isolate the cause; a fp16-versus-bf16 comparison and a per-layer error analysis would be the next step.
- **The 16 longest assays are excluded, and not at random.** Assays beyond 1022 residues exceed ESM-2’s context and were skipped rather than truncated, since a truncation window is a scoring-protocol change that would confound the comparison. The excluded set is 50% low-MSA-depth against 13.9% of those retained and contains no stability assays, so it is a biased sample; post-stratification (Section 3.6) puts the resulting inflation at about +0.004. Configuration behaviour above *L* = 934 is unmeasured.
- **Absolute correlations are convention-dependent.** Our flat mean is the most favourable of four defensible aggregations, spanning 0.019. Comparisons against published ESM-2 numbers must match the convention; the configuration *differences* are stable across all four.
- **Only two results survive multiple-comparison correction.** Seventeen comparisons against fp32 are reported — five configurations at three scales, plus the two W8A8 remedies — against a Bonferroni threshold of 0.05*/*17 = 0.0029. Only int8 dyn at 650M (*p* = 0.0003) and the 3B–650M interaction contrast for int4 wo (*p* = 0.0007) clear it. The individual int4 wo effects at 650M and 3B are nominal only (*p* = 0.0102, 0.0104), and we state them as such rather than as established effects.
- **Dominance is stated on benchmark means.** The Pareto surface in Table 19 uses mean *ρ*, the statistic this paper otherwise argues against selecting on. It is the right basis for a general recommendation and the wrong one for a specific target; a practitioner should run the per-target screen regardless of what the frontier says.
- **The SNR screen is validated in-sample.** Its thresholds are read off the same 3015 combinations they are evaluated on, so the 0% damage rate above SNR = 8 is an in-sample figure and the cut-point is not calibrated on held-out data. It should be treated as a heuristic with a demonstrated direction, not as a guaranteed bound.
- **The asymmetric repair is measured at one scale.** The W8A8 comparison is 3B only; we did not re-run it at 650M or 15B, so the repair is established where the failure was and not shown to be general.
- **The bulk-embedding workload is measured at one scale.** Accuracy, memory and DMS speed are reported at 650M, 3B and 15B, but bulk throughput only at 3B. The claim that the two workloads prefer opposite configurations, and the finding that the conflict dissolves by 15B, are therefore asymmetric in their evidence: the DMS side spans three scales and the embedding side does not. We would not assume the embedding-side ranking is scale-invariant merely because we did not measure it.
- **The signal threshold is arbitrary.** Conditioning the tail on *ρ*_fp32_ *>* 0.3 is a judgement call. We report the sensitivity across cuts from 0.0 to 0.4; the 3B int8 dyn result is invariant, the 650M int4 wo result is not.
- **Bootstrap resolution.** Benchmark-level tests use *R* = 20,000, so the smallest reportable two-sided *p* is 10^−4^; single-assay tests use *R* = 2000 and floor at 0.001. Values at either floor should be read as upper bounds rather than as point estimates.
- **Low-signal assays resolve nothing individually.** Where *ρ*_expt_ ≈ 0.3 the model’s own error dominates and configuration differences fall below the noise floor; such assays contribute to the benchmark mean but carry little information alone.
- **Sequence-length coverage.** Bulk-throughput measurements used sequences of 512–1024 residues. Beyond *L* ≈ 2000 the *O*(*L*^2^) attention term dominates peak memory and quantization’s share of the memory benefit shrinks correspondingly. A representative FASTA from the target workload would settle this.
- **Determinism is established, not assumed, but only within one stack.** Repeating the benchmark reproduces every score bitwise (Section 3.8), so no seed or ensemble over run variation is needed. This says nothing about stability across GPU architectures or torch/torchao versions, which we did not test and which a reader porting these numbers should not assume.
- **The QLoRA track is unbuilt.** int4 wo at 1.61 GB is the natural base, and fine-tuning is the one setting where quantization buys capability rather than efficiency — full 3B fine-tuning requires roughly 45 GB of optimizer state alone.

## 8 Conclusions

### What to run

Choose the model scale first.

- **For variant-effect scoring, use ESM2-650M.** It is statistically indistinguishable from 3B and 15B on ground truth, and every non-dominated point on the accuracy/memory/speed surface is a 650M configuration. Quantizing a larger model to save memory is strictly worse than not using the larger model.
- **Bulk embedding extraction:** fp8 dyn **with** max-autotune. Best on all three axes at once at 3B — 1.15× bf16 throughput, peak memory 12.63 → 7.24 GB, accuracy indistinguishable from bf16. Needs FP8 hardware (sm 90); on A100, int8 wo.
- **Variant-effect scoring up to 3B:** bf16**, eager, uncompiled.** Fastest at every problem size from *L* = 37 to 934 and 6.2× faster than fp32 over the full benchmark. Quantization is pure overhead here and compilation is actively harmful.
- **Variant-effect scoring at 15B:** fp8 dyn. It matches bf16 throughput (1.01×) at half the peak memory (14.6 vs 28.6 GB) with no measurable accuracy cost. The recommendation inverts with scale.
- int8 dyn **is repairable;** int4 wo **is not, here.** int8 dyn’s −0.37 collapse is a defect of its default *symmetric* activation scaling: switching to asymmetric — one keyword, no calibration — removes all six damaged assays and cuts the worst case to −0.04, and SmoothQuant does the same at higher cost. The accuracy repair transfers to bulk extraction (embedding cosine 0.9896 → 0.9995); its *speed* advantage does not, reversing from 1.95× symmetric on DMS to 0.86× on bulk. Neither configuration is fast enough to displace bf16 for variant-effect scoring, but the blanket advice to avoid W8A8 on ESM-2 would have been wrong.

### What the evidence says about evidence

Four findings generalise past ESM-2, and each contradicts a common practice.

1. **Select on worst-case drift, not on benchmark means.** At no scale does any configuration move mean correlation by more than 0.007 — the statistic most likely to be published, and the one least able to distinguish these configurations. int8 dyn passes the mean test at 3B (*p* = 0.34) and takes one assay from *ρ* = 0.591 to 0.223. On means, int8 dyn and bf16 differ by 0.002; on worst case, by 0.36 — a factor of 180.
2. **Drift from fp32 is a valid safety certificate and an invalid quality ranking.** Over 3015 assay/configuration pairs it predicts the *magnitude* of ground-truth change (*r* = 0.56–0.81) but not its *direction*, whose correlation changes sign across model scales. A configuration close to fp32 is safe; one far from fp32 is merely unpredictable, not worse. bf16 and int8 wo never exceed 0.018 Spearman on any of 402 assay/model pairs up to 3B and are safe on that basis alone — cheap to verify on a target, with no experimental labels required (Algorithm 3).
3. **Conclusions are bounded by model scale, not only by sample size.** The workload conflict that motivates half this paper holds at 650M and 3B and dissolves at 15B. The int4 wo effect differs significantly between 650M and 3B (*p* = 0.0007) without being monotone in scale, so it is established that a one-scale result does not transfer, and not established what the effect is at any single scale. Even the stratification by MSA depth replicates at two scales and breaks at the third. A result measured at one scale should be reported as a result at that scale.
4. **Evidence scale is not a matter of rigour but of conclusions.** This study reached four successive verdicts. A synthetic mutational scan ranked configurations confidently and wrongly. Three real ProteinGym assays overturned it and produced two further generalisations — that quantization does not systematically degrade accuracy, and that checkpoint choice dominates precision choice — both of which the full 201-assay benchmark reversed. That benchmark at two scales then suggested quantization sensitivity grows with model size, which the third scale removed. Every intermediate conclusion was statistically sound on its own data and looked well-supported. The full three-scale benchmark costs 16.2 GPU-hours.

### The general claim

A benchmark mean answers whether a configuration is safe *on average over targets*. Deployment asks whether it is safe on *one* target. These questions have different answers here, and the gap between them is not a detail — it is the difference between 0.002 and 0.36. Wherever a benchmark aggregates over heterogeneous instances and the user faces one instance, we expect the same gap, and we would report the tail alongside the mean as a matter of course.

## 9 Data and Code Availability

All assay data is redistributed from ProteinGym [11], release v1, obtained from the OATML-Markslab/ ProteinGym v1 dataset repository. No new experimental measurements were produced.

The following are released with this work:

- Benchmark harness, scoring driver, and analysis code, including the extraction script that validates every variant’s wild-type residue against target seq before writing.
- **Per-variant predictions** for all 20 model/configuration combinations — six configurations at each of three scales, plus the two W8A8 repairs at 3B — at 2,413,913 scores each, in the canonical assay row order, so that any alternative statistic can be recomputed without access to a GPU.
- P er-assay statistics: 4020 rows in total (1206 at 650M and 15B, 1608 at 3B), giving accuracy, wall-clock, peak memory and fidelity for every assay/configuration pair, together with the aggregated summaries reported in Section 4.

Code, per-assay statistics, and both documents are available at https://github.com/qshao/esm2-quantization.

The per-variant predictions are excluded from that repository on size grounds (≈90 MB compressed) and are regenerated by the batch script it contains; the derived per-assay table, from which every result in this paper is computed, is included in full.

## A Supporting Tables

These four tables support claims made in the main text but are not themselves part of the argument: two describe the measurement setup, and two give the three-assay pilot detail that Section 4.7 supersedes.

**Table 24:**
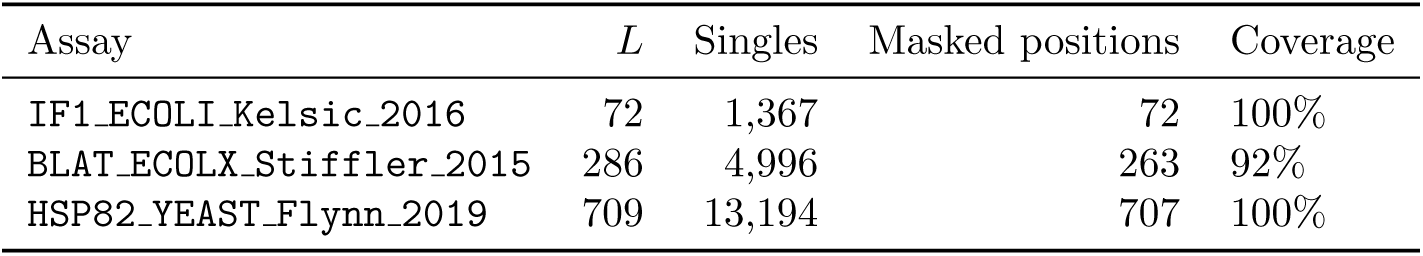
ProteinGym assays used for the three-assay pilot (Sections 4.1–4.6). All 19,557 single-substitution variants had their stated wild-type residue verified against the reference target seq before use; all matched. This check is load-bearing — an off-by-one in position indexing does not raise an error, it silently scores the wrong positions and yields a plausible-looking correlation.

**Table 25:** Hardware and software environment. The glibc 2.17 ceiling is what forces torch 2.6.0 and rules out the prebuilt bitsandbytes GPU wheels, so NF4 was unavailable and is absent from the configuration list.

| Component | Detail |
| --- | --- |
| GPU (primary) | NVIDIA H200, 143 GB, sm_90 (FP8 tensor cores) |
| GPU (secondary) | NVIDIA A100, 80 GB, sm_80 (no FP8) |
| Driver | 550.54.14 |
| PyTorch | 2.6.0+cu124 |
| Attention | SDPA |
| Compile mode | max-autotune-no-cudagraphs |

**Table 26:** Peak device memory during DMS scoring, ESM2-3B, at the three pilot assay lengths. Weights account for approximately 90% of peak even at *L* = 709. Referenced from Section 4.4.

| Config | Weights (GB) | Peak $L=72$ | Peak $L=286$ | Peak $L=709$ |
| --- | --- | --- | --- | --- |
| fp32 | 10.72 | 10.87 | 11.19 | 11.85 |
| bf16 | 5.29 | 5.38 | 5.55 | 5.88 |
| int8_wo | 2.66 | 2.77 | 2.92 | 3.25 |
| int8_dyn | 2.65 | 2.87 | 2.91 | 3.36 |
| fp8_dyn | 2.65 | 2.76 | 2.96 | 3.35 |
| int4_wo | **1.61** | **1.70** | **1.87** | **2.20** |

## B Reproduction

\# `Environment (handles glibc/GCC/cache-quota constraints) source env/activate.sh`

\# `Fetch assay data; verifies variant indexing against target_seq python src/fetch_proteingym.py IF1_ECOLI_Kelsic_2016 \ BLAT_ECOLX_Stiffler_2015 HSP82_YEAST_Flynn_2019`

\# `Bulk throughput + fidelity matrix (compiled)`

**Table 27:**
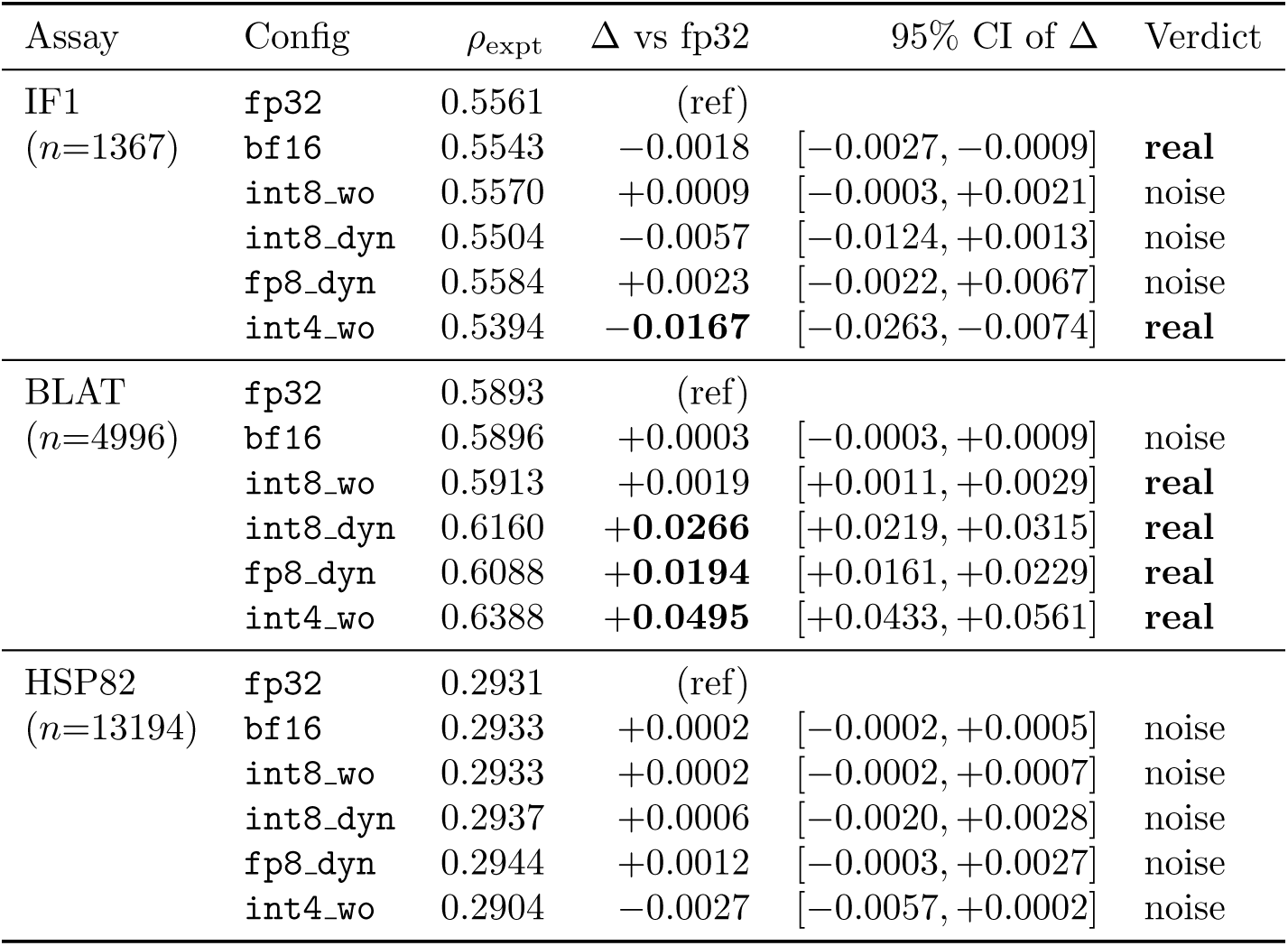
Three-assay pilot: Spearman correlation against experimental measurements, ESM2-3B. Δ is the change from fp32, with a 2000-sample paired bootstrap over variants. “Real” indicates the 95% confidence interval excludes zero. Referenced from Section 4.5; the benchmark-scale replacement is Table 7.

| Assay | Config | $\rho_{\text{expt}}$ | $\Delta$ vs fp32 | 95% CI of $\Delta$ | Verdict |
| --- | --- | --- | --- | --- | --- |
| IF1<br>($n=1367$) | fp32 | 0.5561 | (ref) | | |
| | bf16 | 0.5543 | -0.0018 | $[-0.0027, -0.0009]$ | **real** |
| | int8_wo | 0.5570 | +0.0009 | $[-0.0003, +0.0021]$ | noise |
| | int8_dyn | 0.5504 | -0.0057 | $[-0.0124, +0.0013]$ | noise |
| | fp8_dyn | 0.5584 | +0.0023 | $[-0.0022, +0.0067]$ | noise |
| | int4_wo | 0.5394 | **-0.0167** | $[-0.0263, -0.0074]$ | **real** |
| BLAT<br>($n=4996$) | fp32 | 0.5893 | (ref) | | |
| | bf16 | 0.5896 | +0.0003 | $[-0.0003, +0.0009]$ | noise |
| | int8_wo | 0.5913 | +0.0019 | $[+0.0011, +0.0029]$ | **real** |
| | int8_dyn | 0.6160 | **+0.0266** | $[+0.0219, +0.0315]$ | **real** |
| | fp8_dyn | 0.6088 | **+0.0194** | $[+0.0161, +0.0229]$ | **real** |
| | int4_wo | 0.6388 | **+0.0495** | $[+0.0433, +0.0561]$ | **real** |
| HSP82<br>($n=13194$) | fp32 | 0.2931 | (ref) | | |
| | bf16 | 0.2933 | +0.0002 | $[-0.0002, +0.0005]$ | noise |
| | int8_wo | 0.2933 | +0.0002 | $[-0.0002, +0.0007]$ | noise |
| | int8_dyn | 0.2937 | +0.0006 | $[-0.0020, +0.0028]$ | noise |
| | fp8_dyn | 0.2944 | +0.0012 | $[-0.0003, +0.0027]$ | noise |
| | int4_wo | 0.2904 | -0.0027 | $[-0.0057, +0.0002]$ | noise |

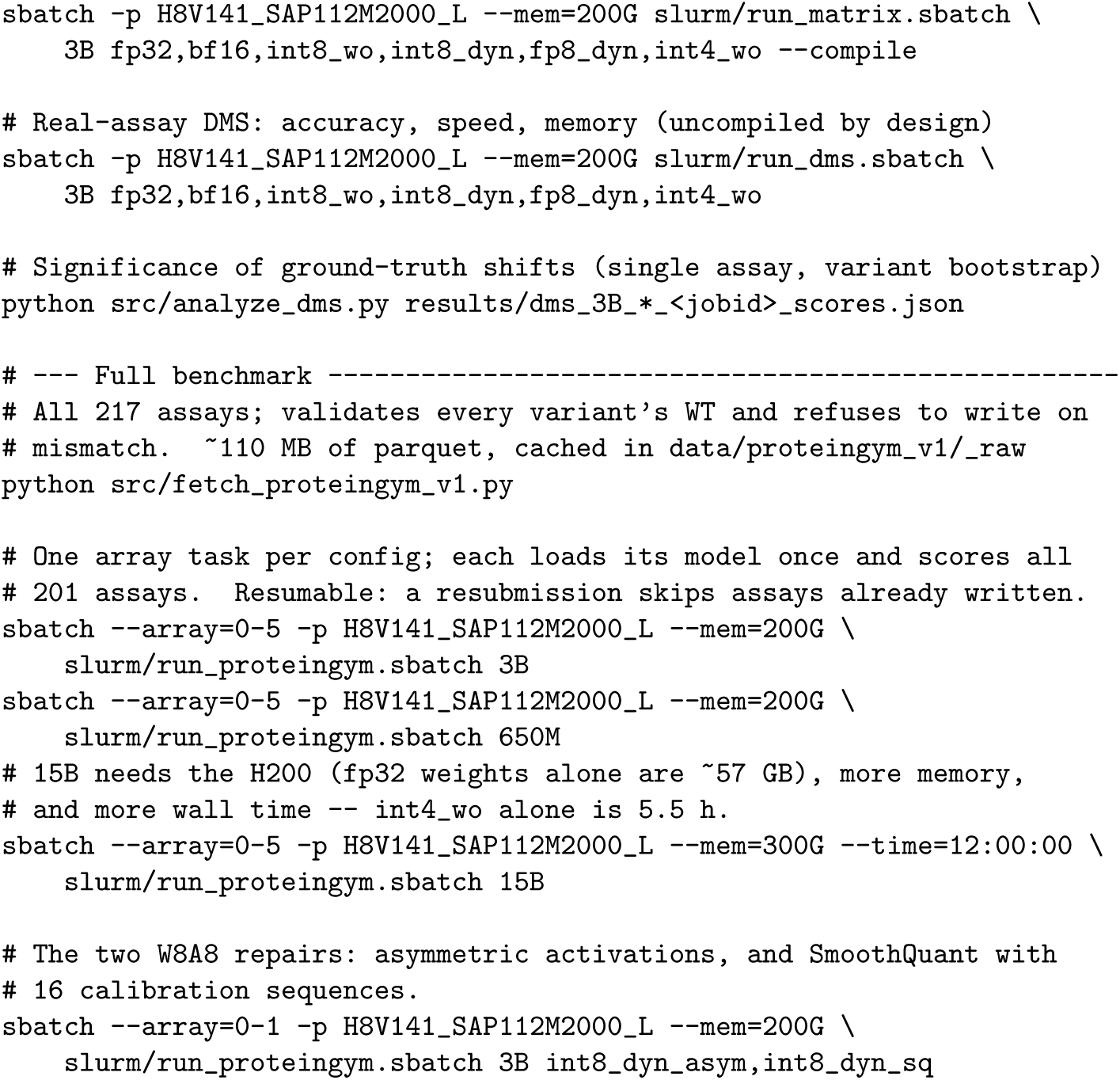
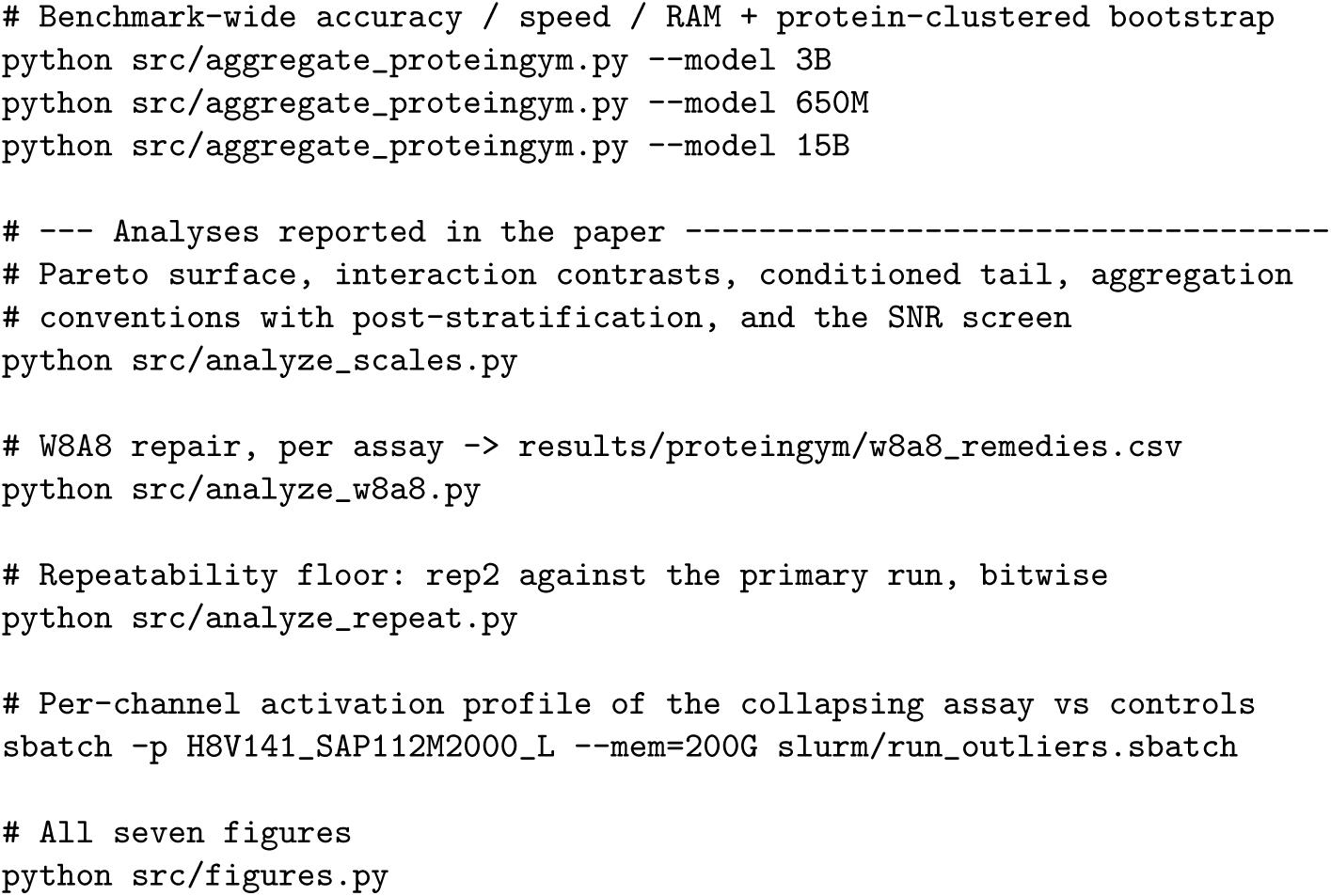

Result artifacts referenced in this report:

- Throughput matrices (SLURM jobs 5393462, 5393520, 5393524, 5393554): results/matrix 3B <jobid>.json
- ProteinGym, 3B, with per-variant scores (job 5394386): results/dms 3B <assay> 5394386.json
- ProteinGym, 650M (job 5394316): results/dms 650M <assay> 5394316.json
- Full benchmark, per-variant scores, 3B (array 5395036) and 650M (array 5395037): results/proteingym/<model> <config>.jsonl.gz
- Full-benchmark analysis: results/proteingym/summary <model>.json andsummary <model> per assay.csv (one row per assay/con_guration: 1206 at 650M and 15B, 1608 at 3B)

